# Drought effects of annual and long-term temperature and precipitation on mortality risk for 9 common European tree species

**DOI:** 10.1101/2022.11.10.515913

**Authors:** Matthias Neumair, Donna P. Ankerst, Nenad Potočić, Volkmar Timmermann, Mladen Ognjenović, Susanne Brandl, Wolfgang Falk

## Abstract

Risk factors for natural tree mortality in managed forests, excluding wind and snow induced breakage, fires and thinning, can be difficult to identify due to correlated confounders of long- and short-term weather patterns with tree age. This study quantified the association of annual and long-term 30-year average temperature and precipitation effects on individual tree death across Europe from 2011 to 2020 for European beech, sessile and pedunculate oak, silver birch, black pine, Austrian oak, Scots pine, European hornbeam, and Norway spruce. For each species, logistic regression approaches for predicting annual mortality evaluated the influence of age, exposition and weather effects on individual tree death, while accounting for multi-collinearity of risk factors. For all species except sessile oak, higher 30-year-temperature averages were associated with higher odds of tree mortality. Effect size of other risk factors varied among species, with similar weather associations between Austrian and sessile oak on the one hand, and Scots pine, Norway spruce and pedunculate oak on the other hand. In particular, warmer winters reduced mortality for silver birch, sessile and Austrian oaks, while having the opposite association for the other species. Sessile oak was most robust against drought effects and could serve as an important tree species under climate change scenarios.

## 1. Introduction

Mortality as premature tree death is a ubiquitous phenomena in ecosystems in space and time that can manifest as a diffuse background process, as self-thinning in stands, or due to pulse events that include phenomena, such as heat waves, droughts, fires, floods, windstorms, ice storms, snow, and pest outbreaks. The frequency and magnitude of the latter is expected to change because of human behaviors interacting with climate and land-use change, as well as species invasions (Jentsch and White 2019). In addition, background tree mortality is likely to be affected by gradual climate changes, such as increasing temperature and decreasing water availability (Taccoen et al. 2019). Forest management with a tendency towards even-aged monocultures on sites outside the natural range of species, as in the case of Norway spruce in Europe (Caudullo et al 2016), is another factor increasing mortality risk (Brandl et al. 2020). Mortality of dominant trees in stands results in economic loss (Hanewinkel et al. 2013) and reduces ecosystem services to various degrees (Thom and Seidl 2016; Mueller et al. 2005).

Mortality events can either happen fast (as a result of storms, fire, bark beetle outbreaks) or rather slow, through various decline processes (most often interactions including climate, insects, or pathogens like root rot or blue stain fungi). Previous studies have found associations between tree mortality and climate (Brandl et al. 2020), weather (Neumann et al. 2017), soil (Maringer et al. 2021), stand parameters (Maringer et al. 2021; Taccoen et al. 2021), and insects and diseases (Anderegg et al. 2015). However, tree mortality is a complex process (Anderegg et al. 2015; Manion 1991) that is difficult to decipher due to competing and largely unobserved mechanisms operating over large climatic gradients. For example, weather has both direct and indirect effects on tree vitality, e.g. via its impact on insect populations (Anderegg et al. 2015).

Climate change influences many of the processes related with disturbances and tree mortality (Romeiro et al. 2022). Warmer winters influence growth in temperature limited regions, but in boreal or temperate forests also increase risks for frost damage (missing snow cover), insect attacks (reproduction and survival), wildfire, and root rot (Romeiro et al. 2022). Rising temperatures have major impacts on drought severity due to vapor pressure deficit leading to climate change-type drought (Carnicer et al. 2011), hot or hotter drought (Allen et al. 2015). Although drought has historically shaped European forests (e.g., Przybylak et al. 2020), steep temperature rises are likely to increase frequency and magnitude of record events (Allen et al. 2015; Park Williams et al. 2013; Forzieri et al. 2021; Senf and Seidl 2021a). These phenomena strike managed forests that are different to natural forests in many ways and have a tendency for lower resilience and robustness due to low complexity (species, genetics, structure, thinning regime).

There are two interrelated physiological mechanisms that are associated with drought induced tree mortality. The first is hydraulic failure through partial or complete loss of xylem function from embolism that therefore inhibits water transport, leading to tissue desiccation, and the second is carbon starvation as a consequence of stomatal closure. Reduced carbon uptake leads to imbalance between carbohydrate demand and supply, resulting in difficulties to meet osmotic, metabolic, and defensive carbon requirements which weakens trees and makes them more vulnerable to biotic and abiotic stresses. In extreme cases, this can lead to mortality by carbon starvation (McDowell and Allen 2015; Adams et al. 2017).

Tree species (and regionally adapted ecotypes) do differ in their tolerances to environmental stress and therefore their ability to withstand changes in climate and climate-induced disturbances (Wang et al. 2012). Choat et al. (2018) state that “plants have limited physiological potential to respond to rapid changes in the environment” due to the fine balance of carbon gain and risk of hydraulic failure. So, despite a general adaptation to environmental conditions, e.g. in life expectancy, growth rates, root to shoot allocation patterns, leaf area index and, leaf phenology, stomatal control or water use efficiency that reflect their differing distribution, strong changes in climate have the potential to harm trees. Whereas xylem hydraulic failure seems to be ubiquitous across multiple tree taxa at drought induced mortality, evidence supporting carbon starvation is not universal and more common for gymnosperms than angiosperms (Adams et al. 2017). Species differ in their degree to adapt wood anatomical traits (Vander Mijnsbrugge et al. 2020) or e.g. the degree in which they can compensate for leaf damage caused by insects during the spring (e.g. summer shoots of *Q. petraea* and *Q. robur*). Furthermore, tree species suffer from specialized insects or pests, for example Norway spruce is strongly affected by the bark beetle *Ips typographus* (McDowell and Allen 2015). Shallow root system of spruce intensifies drought risk (Caudullo et al. 2016; Netherer et al. 2019). Another example of differences is the consequence of premature leaf shedding, which in the case of beech, with its thin bark, can lead to bark damage and a spiral of decline (Schuldt et al. 2020).

Managed forests in Europe accumulated substantial amounts of biomass in the 20th century. The changes in forest structure in combination with climate change and other human impacts led to an episode of increasing forest disturbances in recent decades (Senf and Seidl 2021b). In order to fulfill ecosystem services in the future, it is important to increase resilience of managed stands and to do that, understand causes and patterns of mortality. Long-term climate values, such as 30-year temperature or precipitation averages, constitute predisposing factors that may lead to a slow decline and a gradual increase in background mortality. However, an effect on mortality might become only apparent in combination with inciting factors, i.e. current or very recent annual climate anomalies. To capture these effects this study investigated data series until 2020 as the 2018 drought and heat wave in central and eastern Europe was described to be accompanied with large scale mortality and lag effects (Senf and Seidl 2021a; Schuldt et al. 2020; Brun et al. 2020). Revealing the links between long-term climate data, short-term weather anomalies and mortality could help improve management. This study focuses on long-term 30-year average climate (temperature, precipitation) and yearly weather anomaly-induced mortality patterns in Europe as these are expected to increase dramatically (Forzieri et al. 2021; Senf and Seidl 2021a).

Because mortality is a rare event in forests, short-term or small-scale studies lack power for identifying key management and climatic indicators. The pan-European ICP Forests crown condition data set covers a large climatic gradient and therefore provides a rich information source for mortality modeling studies (Brandl et al. 2020; Neumann et al. 2017). Therefore, this data set was used in this study despite a limited information depth on stand characteristics and soil information, both factors known to be related with mortality (Maringer et al. 2021; Taccoen et al. 2021). Time series was limited to 2011-2020 due to the introduction of the removal code in the data in 2011 as well as to control for common long-term average effects in the analysis of most recent and relevant data for contemporary forests.

The hypotheses of this study are: (i) Individual tree mortality is driven by both annual anomalies and long-term 30-year average temperature and precipitation effects, interacting with stand variables and species. (ii) The effects of drought in Europe can be witnessed via the effects of temperature and precipitation on tree mortality in the drought year and the following years. (iii) Although heterogeneous by species, risk mortality profiles among species that grow at similar sites may exhibit similarities.

## 2. Results

In total, 746,478 tree-year observations were recorded among nine species during the years 2011-2020 (Table 1).

**Table 1.** European tree species present in the study with data on tree-year observations in the period 2011-2020.

| Scientific name | Common name | Alive (n) | Dead (n) |
| --- | --- | --- | --- |
| *Fagus sylvatica* | European beech | 147930 | 419 |
| *Pinus nigra* | black pine | 45102 | 132 |
| *Picea abies* | Norway spruce | 156740 | 1399 |
| *Pinus sylvestris* | Scots pine | 225722 | 953 |
| *Betula pendula* | silver birch | 21549 | 145 |
| *Quercus cerris* | Austrian oak | 28536 | 101 |
| *Carpinus betulus* | European hornbeam | 17550 | 103 |
| *Quercus robur* | pedunculate oak | 47945 | 277 |
| *Quercus petraea* | sessile oak | 51643 | 232 |
| Total |  | 742717 | 3761 |

Annual rates of mortality for the nine species, shown in Figure 1, vary from near 0% to above 2.5% and show peaks after the 2013, 2015 and 2018 drought periods in Central Europe. Norway spruce and silver birch exhibited the highest values of mortality exceeding 1.5% in 2019 and respectively, 2020. Figure S1 and S2 overlay the trends in mortality according to annual average temperature and precipitation variables, showing how increases in temperature or decreases in precipitation could induce increases in mortality for all species; Figures 3 and 5 to 12 show the corresponding plots by individual species. Significant associations between mortality and weather characteristics are summarized in Figure 2, and show heterogeneity of effects across the 9 species. Multivariate risk models across species performed relatively well for predicting mortality, with all areas under the receiver operating characteristic curves (AUCs) exceeding 67%.

**Fig 1.**
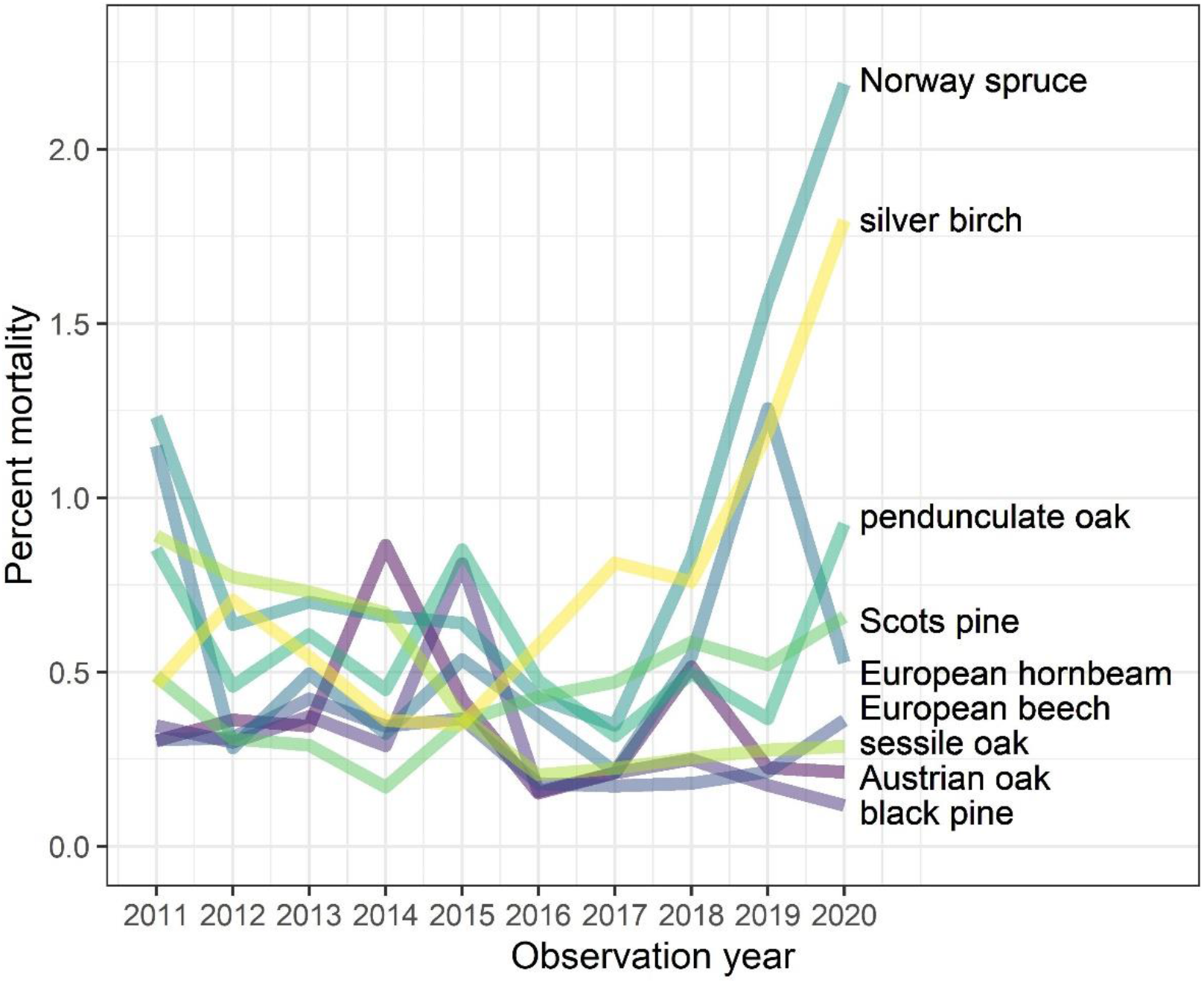
Annual mortality by species from 2011 to 2020

**Fig 2.**
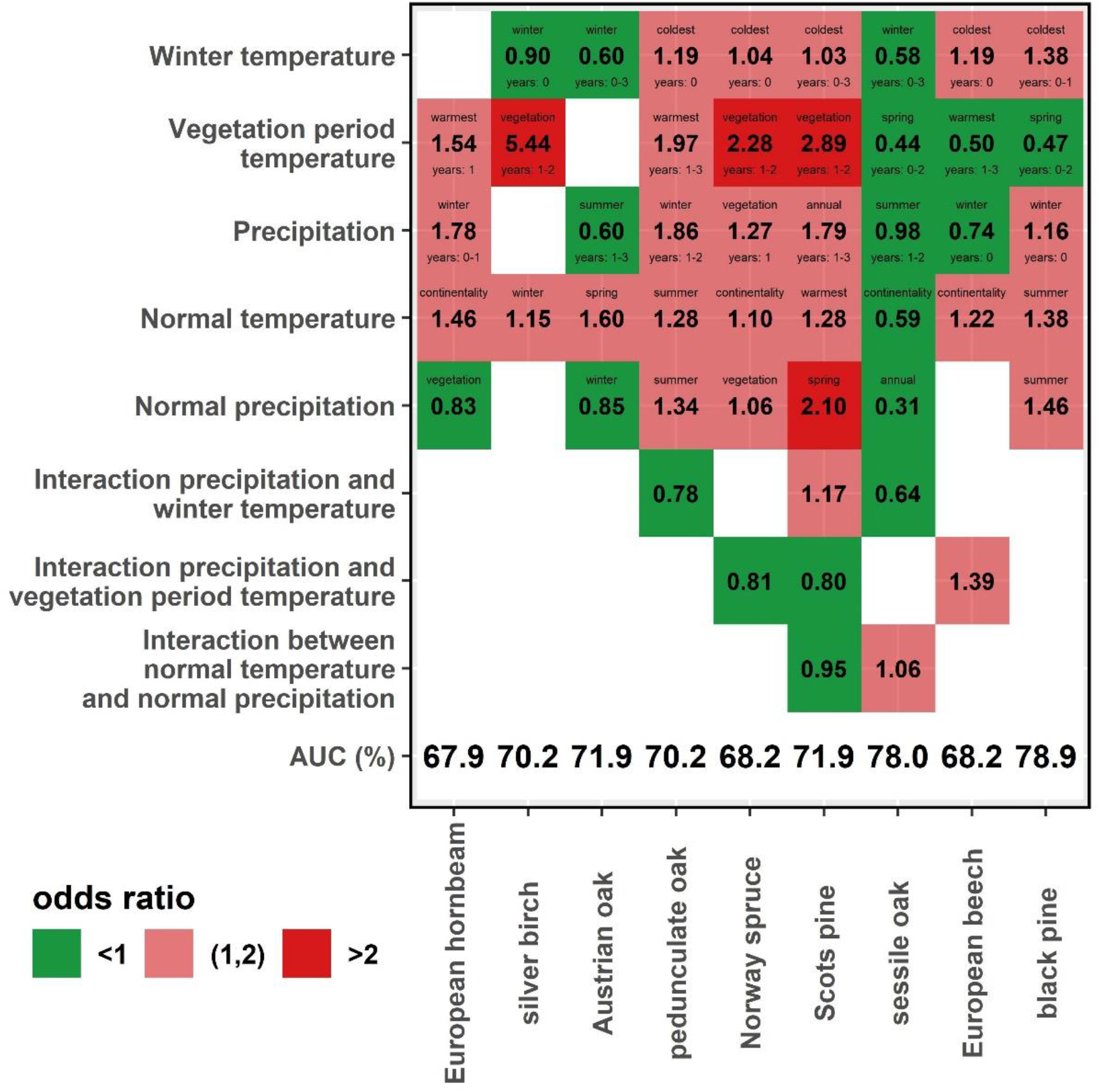
Odds ratio (OR) estimates based on stand and weather characteristics of all species by variable group. OR’s above 1 indicate that the increasing values of the risk factor increases the risk of tree death, while OR’s below 1 indicate increasing values of the risk factors decrease the risk of tree death. OR’s equal to 1 indicate the risk factor has no impact on tree death. Normal indicates the average of the annual respective weather variable measurements during the 30-year period 1981-2010. For variables without this prefix the anomaly is calculated by subtracting the climate normal from the seasonal value at plot level. Red indicates an increase in tree death odds, green a decrease. Area underneath the receiver operating characteristic curve (AUC) indicates discrimination of a risk model for predicting the outcome of tree death, with higher values indicating better discrimination of trees that experienced death compared to those that did not. All effects were significant (p < 0.05) except winter temperature in Scots pine (OR=1.03) and precipitation for sessile oak (OR=0.98). The selected variable within the variable group (Table 2) is indicated with abbreviations beside the averaged years for annual variables. Further model details can be found in Tables S1-S9

**Fig 3.**
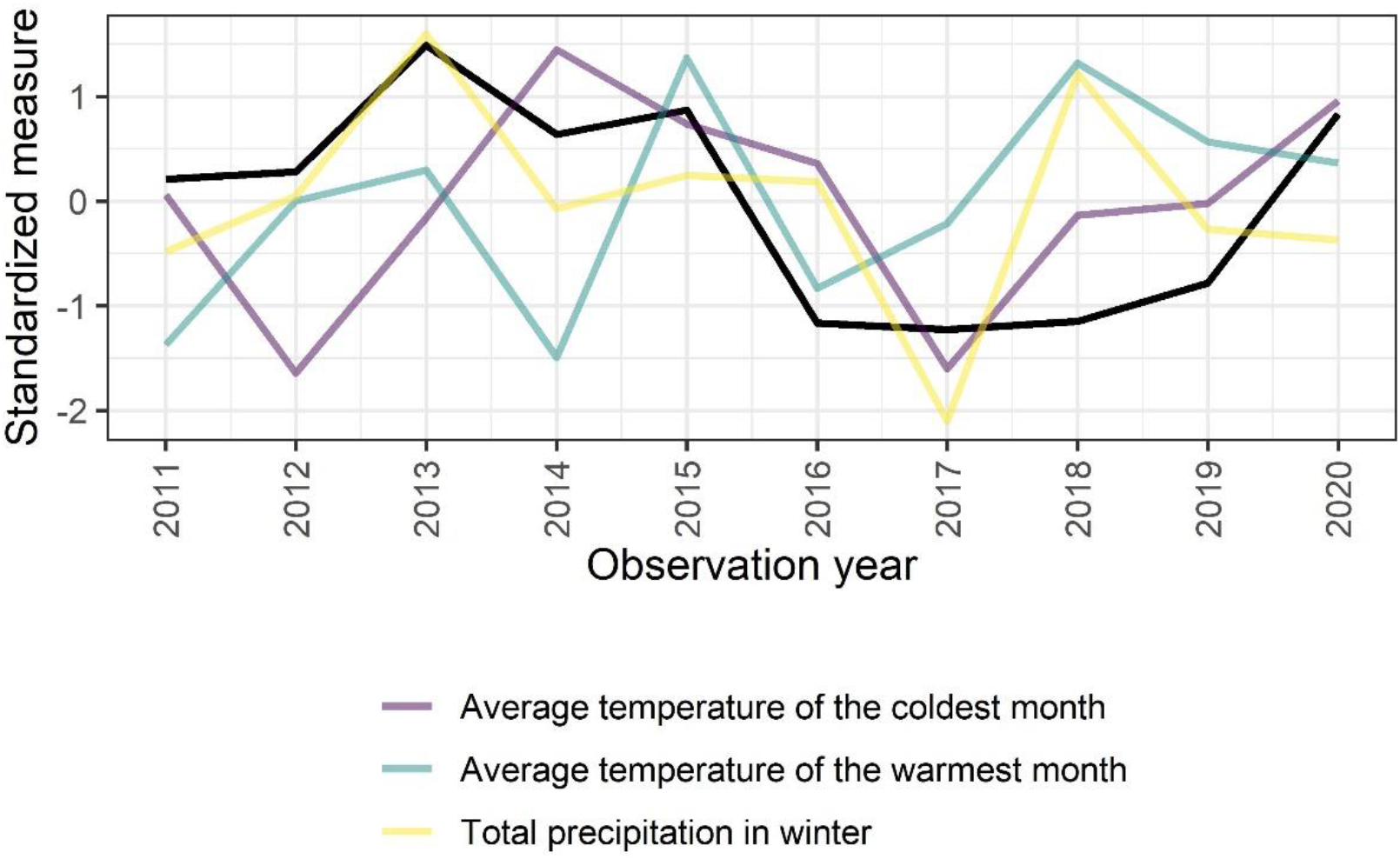
Annual values of weather variables and mortality (black line) for European beech. All variables have been standardized by subtraction of their means and divided by their standard deviations. Only variables from the final model in the pooled analysis over all years combined are shown

### European beech

Among 148,349 European beech tree-year observations from 2011 to 2020, 419 trees died (Table 1). European beech had lower annual mortality rates than the other species during most years, and was less affected by the droughts of 2013, 2015, and 2018 than the other species (Figure 1). Stand age of more than 120 years increased the odds of tree mortality by 53% compared to the reference age of 41-60 years (odds ratio (OR) 1.53, Table S1). In comparison with flat terrain, eastern exposition increased the odds of tree mortality by 58%, whereas south-western exposition decreased it by 48%. As a climatic long-term effect, plots with a 1°C higher continentality, defined as the average of the difference between the warmest and coldest month average temperature over the 30 years from 1981 to 2010, experienced increased odds for tree mortality by 22%. Additionally, a 1°C increase in the average temperature of the coldest month of the current winter increased the odds of tree mortality by 19%. This effect was confirmed by the results shown in Figure 3, where with the exception of 2012-13, years with higher average temperature of the coldest month also had increased mortality. This can be seen especially in the drought year 2015. A 50mm increase in winter precipitation of the current year decreased the odds of tree mortality by 26%. This association is seen in Figure 3 from 2018 onward where winter precipitation decreased and mortality increased. A 1°C increase in the average temperature of the warmest month aggregated over the three previous years decreased the odds of tree mortality by 50%. This can be seen especially for the years 2016 to 2019 with less mortality following the temperature peaks in 2015 and 2018 (Figure 3). Combined, an increase in precipitation sum of winter and warmest month mean temperature led to 39% higher odds of tree mortality. The resulting model led to the evaluations shown in Figure 4, indicating higher mortality risks in northern and east-central Europe, with lower risks further west and in the south, depending on the year in question.

**Table 2.** Weather characteristics. Top variables were available for every year from 2010 to 2020; 30-year averages were calculated during 1981-2010 and remained fixed for every year.

| Variable | Group |
| --- | --- |
| Average temperature of the coldest month (°C) | Winter temperature |
| Average temperature in winter (Dec.(previous year) - Feb.) (°C) | Winter temperature |
| Average temperature of the warmest month (°C) | Vegetation period temperature |
| Average temperature in spring (Mar. - May) (°C) | Vegetation period temperature |
| Average temperature in summer (Jun. - Aug.) (°C) | Vegetation period temperature |
| Average temperature in vegetation period (Mai - Sep.) (°C) | Vegetation period temperature |
| Total precipitation in winter (mm) | Precipitation |
| Total precipitation in spring (mm) | Precipitation |
| Total precipitation in summer (mm) | Precipitation |
| Annual total precipitation (mm) | Precipitation |
| Total precipitation in vegetation period (mm) | Precipitation |
| 30-year normal of average temperature of the coldest month (°C) | Normal temperature |
| 30-year normal of average temperature of the warmest month (°C) | Normal temperature |
| 30-year normal of average temperature in winter (°C) | Normal temperature |
| 30-year normal of average temperature in spring (°C) | Normal temperature |
| 30-year normal of average temperature in summer (°C) | Normal temperature |
| 30-year normal of average temperature in vegetation period (°C) | Normal temperature |
| 30-year normal of temperature difference between average temperature of the warmest and coldest month, or continentality (°C) | Normal temperature |
| 30-year normal of total precipitation in winter (mm) | Normal precipitation |
| 30-year normal of total precipitation in spring (mm) | Normal precipitation |
| 30-year normal of total precipitation in summer (mm) | Normal precipitation |
| 30-year normal of annual total precipitation (mm) | Normal precipitation |
| 30-year normal of total precipitation in vegetation period (mm) | Normal precipitation |

**Fig 4.**
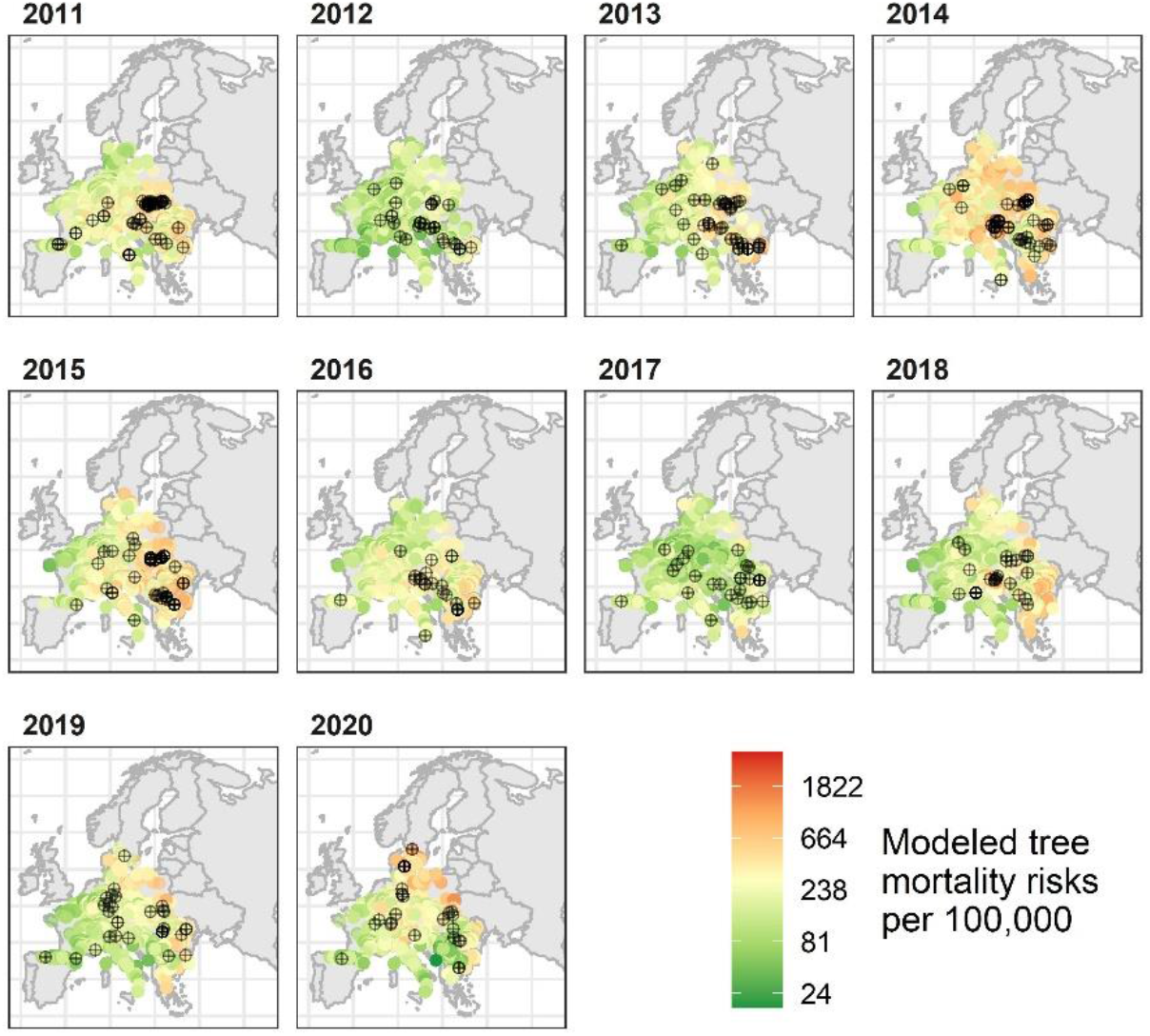
Spatial evaluation of modelled tree mortality for European beech for all analyzed ICP Forests plots on a logarithmic color scale. Black points indicate observations of dead trees (n=419)

### Black pine

Among 451,234 black pine tree-year observations from 2011 to 2020, 132 trees died (Table 1). Black pine had a peak in mortality in 2015 (Figure 1). Stand age of 41-60 years resulted in the highest odds of tree mortality, being 67% to 55% higher than for other age groups (Table S2). For aspect, south, south-east and east exposition increased the mortality odds by up to 392%. As a 30-year climatic long-term effect, plots with more summer precipitation and higher summer temperature had increased tree death odds by 38% and 46%, respectively. Additionally, higher average temperature of the coldest month in the current and previous year increased the odds of tree death by 38%. This association cannot be observed in Figure 5. Higher winter precipitation of the current year increased the odds of tree death by 16% per 50mm, which can be seen especially in the year 2015, but not in 2013 (Figure 5). Higher spring mean temperature of the current year and the two previous years decreased the odds of tree death by 53% per 1°C. This association can be seen in Figure 5 where mortality decreased after the 2018 peak in spring temperature and the mortality peak in 2015 was preceded by lower spring temperatures in the three previous years. The model evaluations indicated higher mortality risks in central Europe, which is the edge of the distribution of this species in Europe (Figure S3).

**Fig 5.**
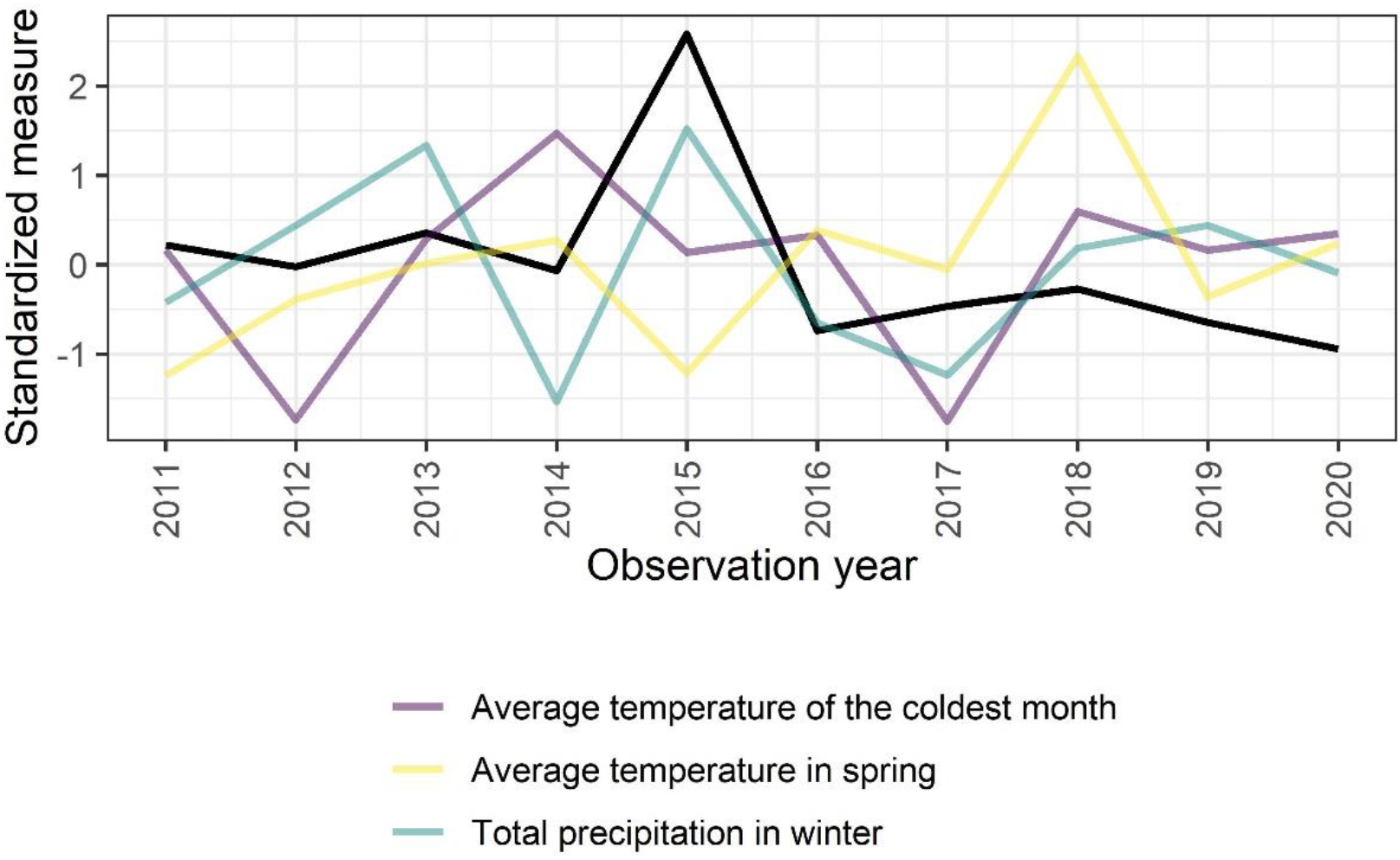
Annual values of weather variables and mortality (black line) for Black pine. All variables have been standardized by subtraction of their means and divided by their standard deviations. Only variables from the final model in the pooled analysis over all years combined are shown

### Norway spruce

Among 156,740 Norway spruce tree-year observations from 2011 to 2020, 1399 trees died (Table 1). Norway spruce had higher mortality rates than most other species during 2011 and from 2018 to 2020 (Figure 1). This effect was strong in central Europe and even southern Sweden (Figure S4). Stand age of 61-80 years resulted in the lowest odds of tree mortality, being 33% lower than for other age groups (Table S3). Norway spruce showed lower odds for south-west, west and north-west aspects (27% - 45%), and higher odds for all other expositions (17% - 70%). As a 30-year climatic long-term effect, plots with more precipitation in the vegetation period and higher continentality had increased tree mortality odds by 6% and 10%, respectively. Additionally, higher average temperature of the coldest month in the current year increased the odds of tree mortality by 4% for 1°C increase, which is driven by the years 2018 to 2020 (Figure 6). During the vegetation period higher average temperature in two previous years and more precipitation in the previous year led to higher tree mortality odds (128% and 27%, respectively), and their interaction to 19% lower tree mortality odds. Figure 6 shows a peak in vegetation period temperature in 2018 accompanied by low precipitation with increased mortality the following years. Model evaluations indicated high mortality risks over the whole of Europe in recent years (Figure S4).

**Fig 6.**
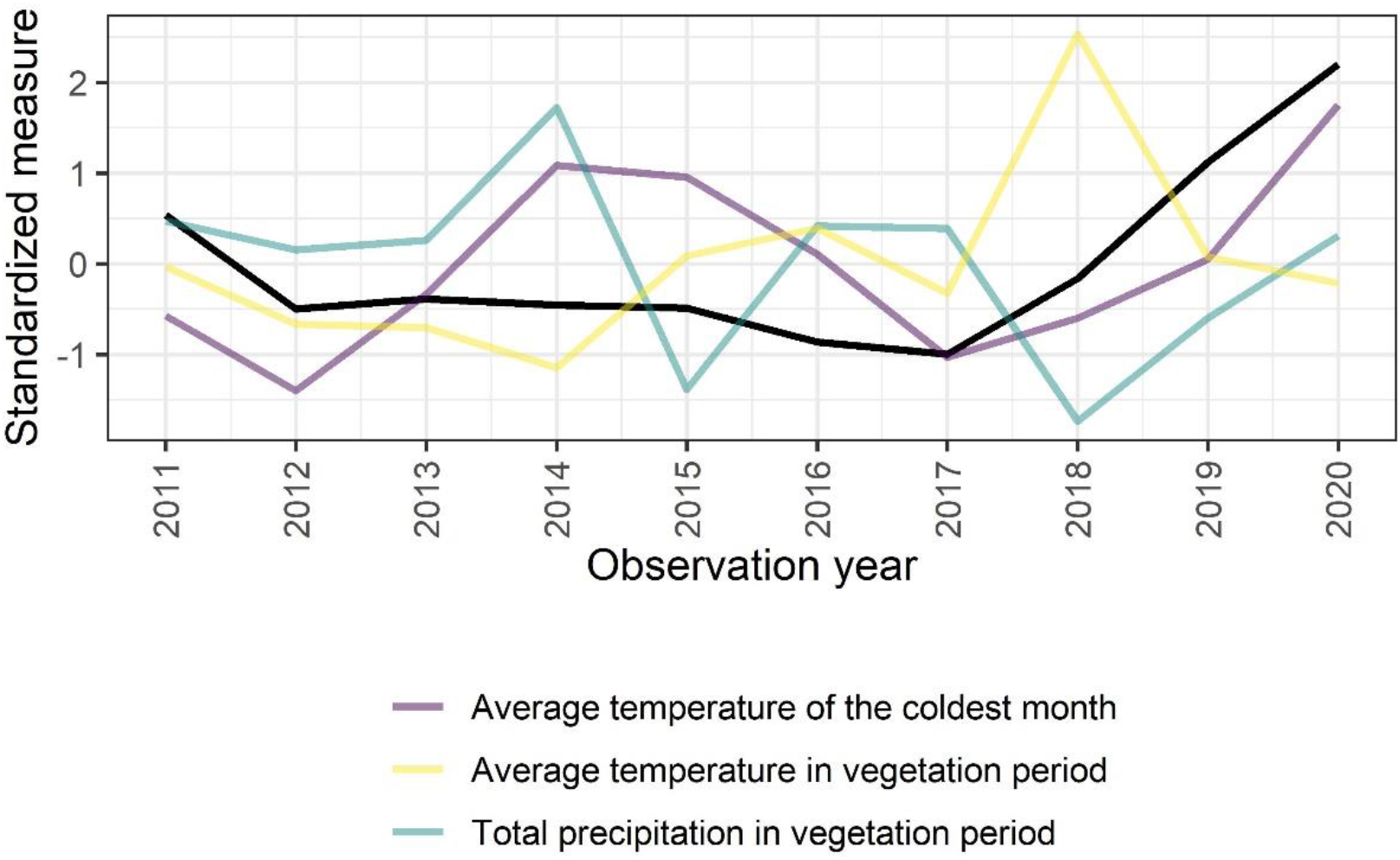
Annual values of weather variables and mortality (black line) for Norway spruce. All variables have been standardized by subtraction of their means and divided by their standard deviations. Only variables from the final model in the pooled analysis over all years combined are shown

### Scots pine

Among 225,722 Scots pine tree-year observations from 2011 to 2020, 953 trees died (Table 1). Scots pine had consistently increasing mortality rates from 2014 to 2020 (Figure 1). Stand age of 41-60 years resulted in the highest odds of tree mortality (Table S4). North-west aspect had 593% higher tree mortality odds compared to flat plots. As a 30-year climatic long-term effect, plots with more spring precipitation and higher average temperature of the warmest month had increased tree mortality odds by 109% and 28%, respectively. However, their interaction reduced tree mortality odds by 5%. Higher average temperature in the vegetation period of two previous years increased the odds of tree mortality by 189%. Figure 7 shows mostly increasing mortality with increasing vegetation period temperature. The temperature peak in 2018 was also seemingly associated with 2020 mortality. Higher average temperature of the coldest month during the three previous years increased the odds of tree mortality by 3%. This association can be seen in Figure 7 where the winter temperature rose from 2012 to 2015 whereas the mortality increased from 2014 to 2020. Also more precipitation the three previous years led to 79% higher tree mortality odds. Figure 7 shows high precipitation from 2012 to 2014, 2016 and 2017 and increasing mortalities from 2014 onward. The interaction between mean vegetation period temperature and the precipitation decreased the tree mortality odds by 20%. The model evaluations indicated no spatial pattern (Figure S5).

**Fig 7.**
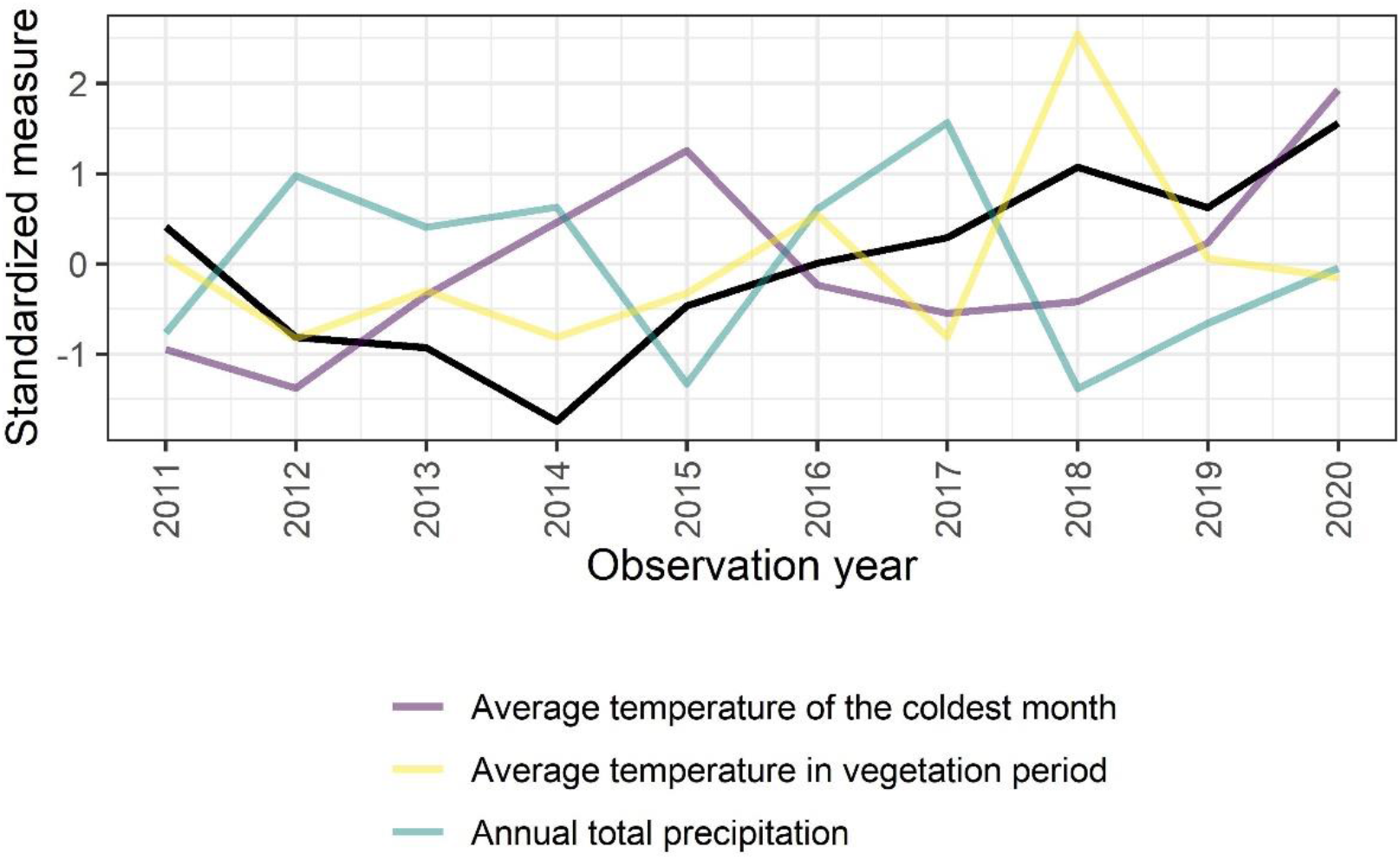
Annual values of weather variables and mortality (black line) for Scots pine. All variables have been standardized by subtraction of their means and divided by their standard deviations. Only variables from the final model in the pooled analysis over all years combined are shown

### Silver birch

Among 21,694 silver birch tree-year observations from 2011 to 2020, 145 trees died (Table 1). Silver birch has had increasing mortality since 2015 (Figure 1). The older tree stands became the higher their odds of tree mortality were (Table S5). As a 30-year long-term effect, plots with higher average winter temperature had increased tree mortality odds by 15%. Higher average vegetation period temperature the two previous years increased the odds of tree mortality by 444%. This association can be seen in Figure 8 where increased vegetation period temperature in 2018 was accompanied by increased mortality in 2019 and 2020. The high temperature in 2018 seems to have had a negative effect until 2020.The average winter temperature in the current year decreased the tree mortality odds by 10%. This association could be seen in the years 2014 to 2016 where the winter temperature increased and the mortality decreased. The model evaluations indicated higher mortality risk in southern Europe with increasing trend over the years (Figure S6).

**Fig 8.**
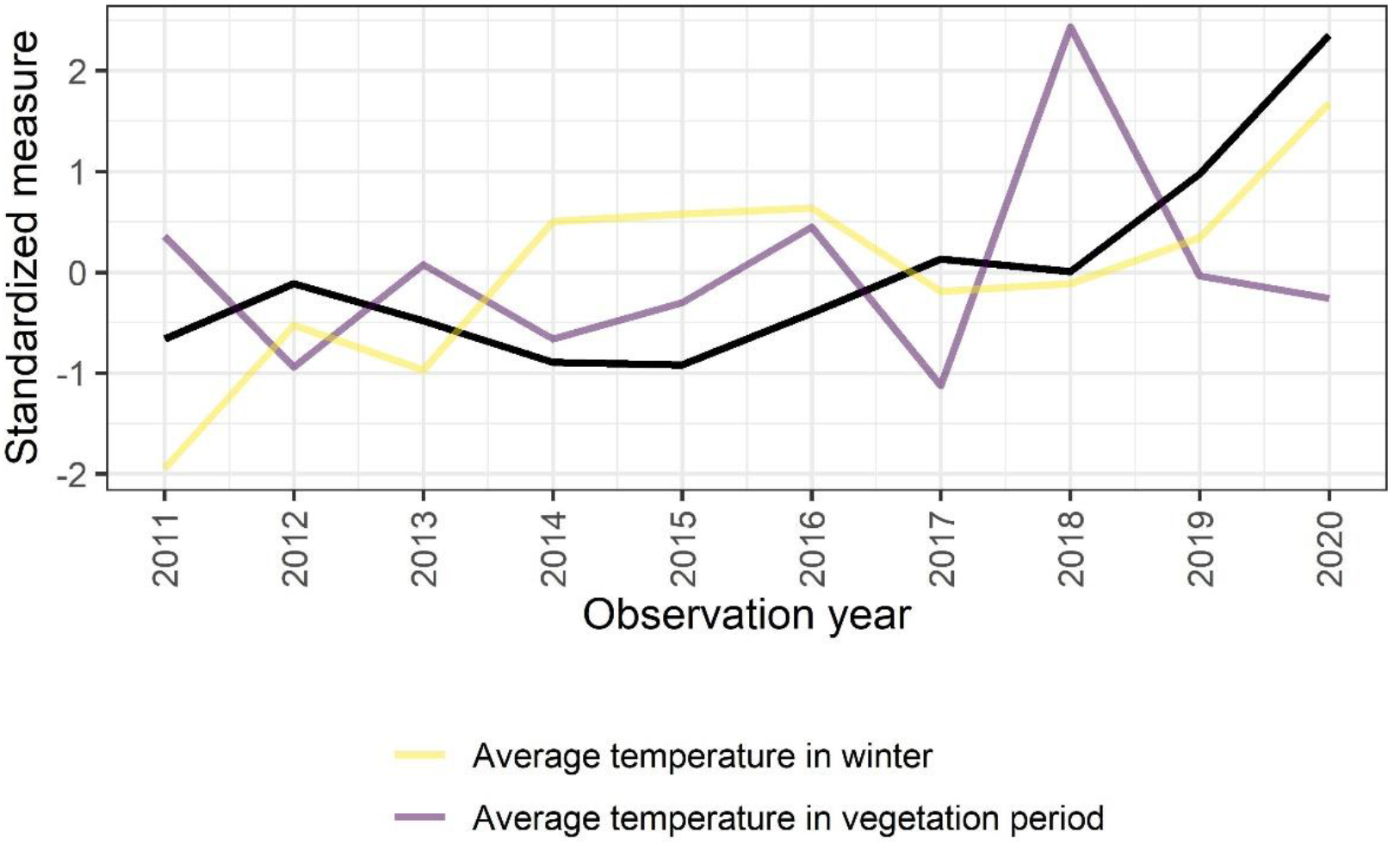
Annual values of weather variables and mortality (black line) for Silver birch. All variables have been standardized by subtraction of their means and divided by their standard deviations. Only variables from the final model in the pooled analysis over all years combined are shown

### Austrian oak

Among 28,637 Austrian oak tree-year observations from 2011 to 2020, all together 101 trees died (Table 1). Austrian oak had the highest mortality in 2014 and a smaller peak in 2018 (Figure 1). This species is spread in south-eastern Europe (Figure S7). 30-year long-term higher average spring temperature increased tree mortality odds by 60% and more winter precipitation decreased tree mortality odds by 15% (Table S6). Higher average winter temperature and also higher summer precipitation during the three previous years decreased the odds of tree mortality by 40%. These opposing trends can be observed for example for 2014 were a temperature and precipitation peak was followed by a mortality decrease during the following years (Figure 9). The model evaluations indicated higher mortality risks in 2013 and 2014 in central Europe (Figure S7).

**Fig 9.**
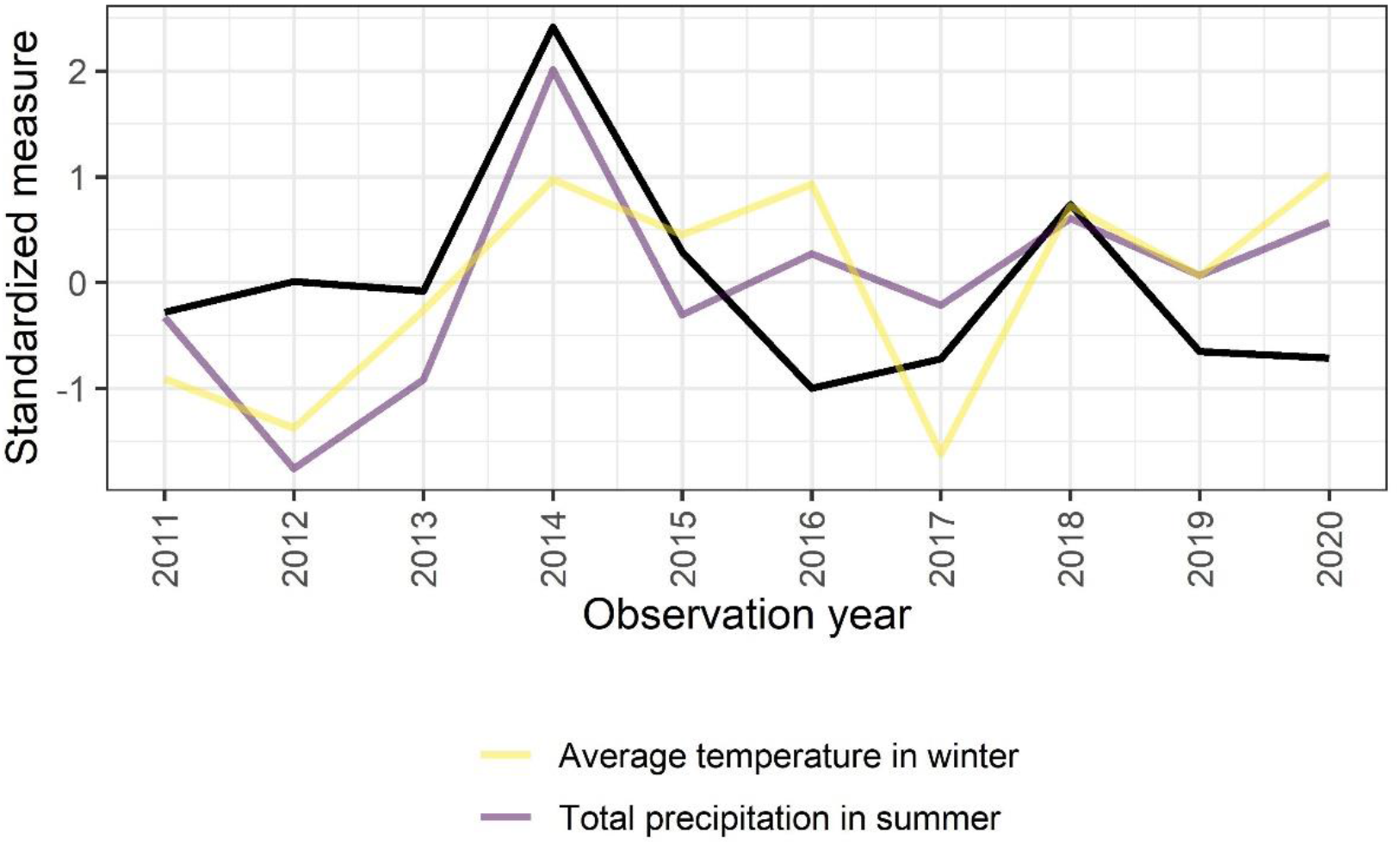
Annual values of weather variables and mortality (black line) for Austrian oak. All variables have been standardized by subtraction of their means and divided by their standard deviations. Only variables from the final model in the pooled analysis over all years combined are shown

### European hornbeam

Among 17,653 European hornbeam tree-year observations from 2011 to 2020, 103 trees died (Table 1). Mortality for European hornbeam was strongly affected in 2011 and 2019 (Figure 1). Tree stands older than 120 years had the highest tree mortality with odds 376% higher than for the reference of 41 - 60 year old stands (Table S7). 30-year long-term higher continentality increased tree death odds by 46% and increased precipitation in the vegetation period decreased tree death odds by 17%. Higher average temperature of the warmest month the previous year and more winter precipitation the current year increased the odds of tree mortality by 54% and 78%, respectively. The temperature association can be seen mostly for the years 2017 to 2019 where an increase in temperature resulted in an increase in mortality the following year (Figure 10). For precipitation, this increasing effect can be seen in all other years except 2011 and 2019. The model evaluations indicated higher risk in eastern Europe (Figure S8).

**Fig 10.**
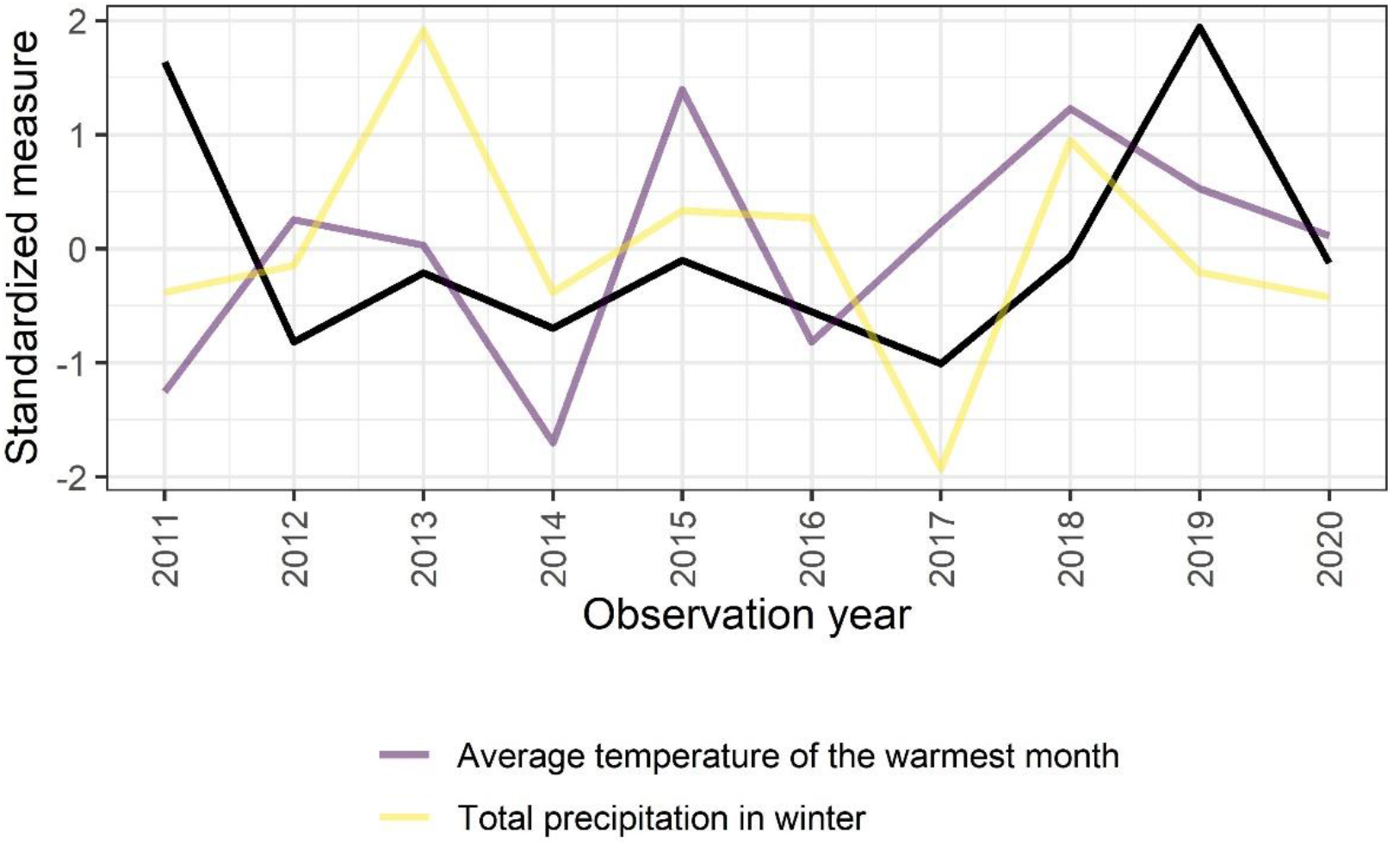
Annual values of weather variables and mortality (black line) for European hornbeam. All variables have been standardized by subtraction of their means and divided by their standard deviations. Only variables from the final model in the pooled analysis over all years combined are shown

### Pedunculate oak

Among 48,222 pedunculate oak tree-year observations from 2011 to 2020, 277 trees died (Table 1). Pedunculate oak exhibited the highest mortality in 2011, 2015 and 2020 with strongest effects in south-eastern parts of central Europe (Figure S9). 30-year long-term higher summer temperature and precipitation increased tree mortality odds by 28% and 34%, respectively. Higher average temperature of the warmest month during the previous three years increased the odds of tree mortality by 97%. Figure 11 shows the immediate association of warmest month temperature for 2015, and the 2018 high temperature association with a two-year delay increase in mortality in 2020. Higher average temperature of the coldest month at the reported year and more winter precipitation during the two previous years increased tree mortality odds by 19% and 86%, respectively, whereas their interaction decreased the mortality odds by 22%. Figure 11 indicates the direct association of coldest month temperature except for 2014, which seemed to influence 2015 mortality. The winter precipitation association seems to be driven by the higher precipitation in 2013 and 2018 with higher mortality in 2015 and 2020.

**Fig 11.**
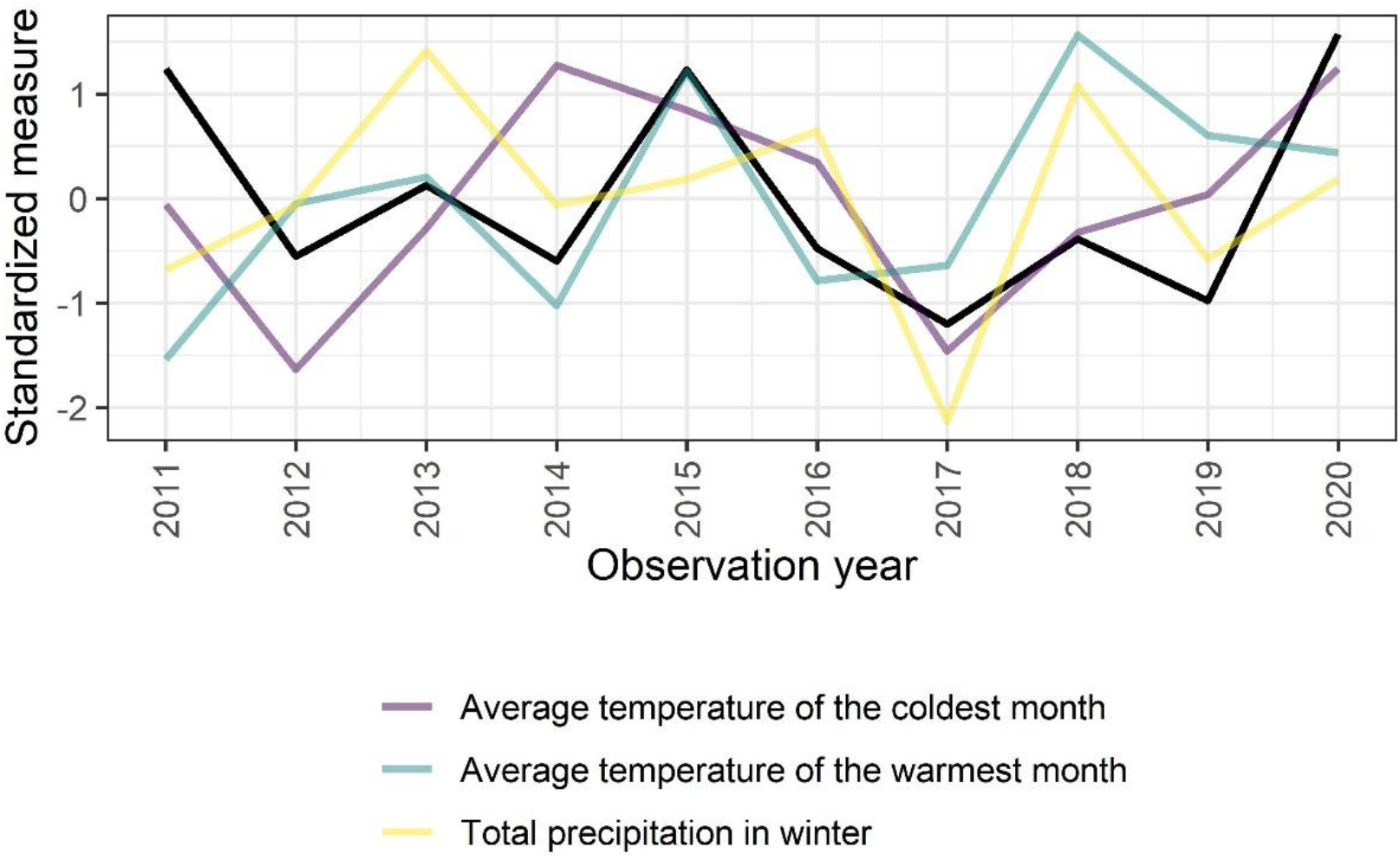
Annual values of weather variables and mortality (black line) for Pedunculated oak. All variables have been standardized by subtraction of their means and divided by their standard deviations. Only variables from the final model in the pooled analysis over all years combined are shown

### Sessile oak

Among 51,875 sessile oak tree-year observations from 2011 to 2020, 232 trees died (Table 1). Sessile oak experienced decreasing mortality from 2011 to 2016 and a slight increase from 2016 to 2020 (Figure 1). Tree stands between 41 and 60 years and plots with north aspects showed highest tree mortality (Table S9). 30-year long-term higher continentality and annual precipitation decreased tree mortality odds by 41% and 69%, respectively, whereas their interaction increased the odds by 6%. Higher average temperature in spring during the previous two years decreased the odds of tree mortality by 56%. Higher average winter temperature during the three previous years and more summer precipitation during the two previous years decreased the mortality odds by 42% and 2%, respectively, and their interaction reduced the odds by a further 36%. These associations can be seen mostly after all three variable peaks in 2014 followed by the mortality decrease the following years (Figure 12). The model evaluations indicated higher mortality risk in central parts of Europe, especially during the years 2013 to 2015 (Figure S10).

**Fig 12.**
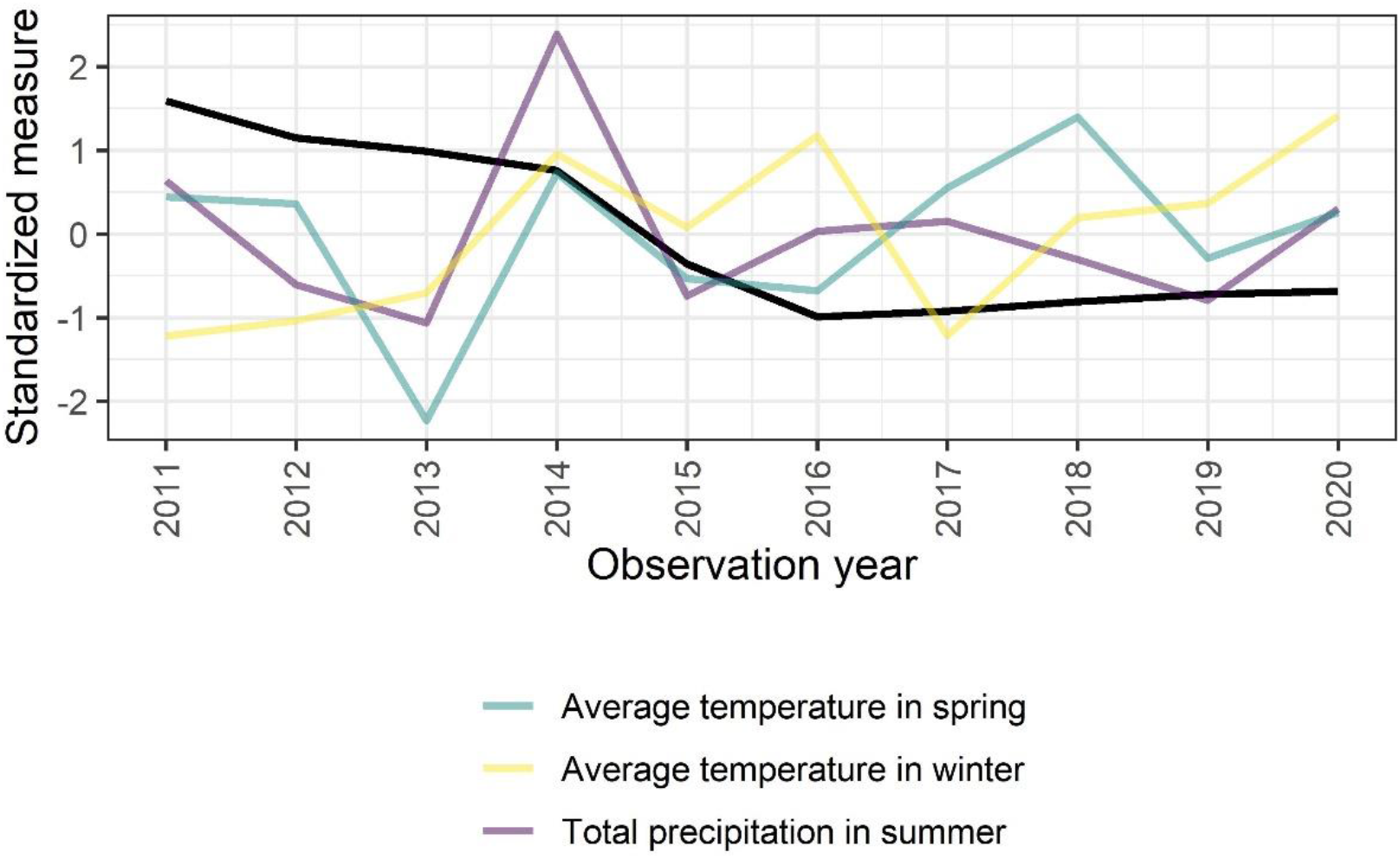
Annual values of weather variables and mortality (black line) for Sessile oak. All variables have been standardized by subtraction of their means and divided by their standard deviations. Only variables from the final model in the pooled analysis over all years combined are shown

## 3. Discussion

The analysis presented here based on annual assessments of common tree species in European forests in the period from 2011 to 2020 showed individual tree mortality to be impacted by long-term climate averages as well as weather factors, in line with previous findings (Brandl et al. 2020; Neumann et al. 2017) forming this study’s first hypothesis.

Long-term climate averages and their change-rate (evolution) over time showed to be of importance for a broad range of species to explain background mortality in France (Taccoen et al. 2019). Likewise, Brandl et al. (2020) found long-term climate variables influencing mortality with higher temperature and lower precipitation increasing mortality risks. In general, higher temperatures within growing seasons are related to higher evaporative demand and therefore might lead to drought stress (Eamus et al. 2013). Additionally, they are favored by pests, as for example, in the case of bark beetles, more generations can fully develop within the season (Marini et al. 2012). Northward and altitudinal spread of forest insects was shown to have a strong temperature dependency, as reviewed in Pureswaran et al. (2018). The high risk of losing climatic niches for many species due to climate warming was also indicated by studies on the predictive distribution of species (Bombi et al. 2017; Dyderski et al. 2018; Thurm et al. 2018) and studies on vegetation samples (Gottfried et al. 2012; Schmidtlein et al. 2013). Brandl et al. (2020) additionally discussed the effect of increased growth and faster development under warmer temperatures, so that trees moved faster along their life’s trajectories and died at younger ages with constant self-thinning lines.

In data of this study, higher temperature normal (winter, spring, summer, warmest month, continentality) was related with higher mortality for all species except sessile oak. Results were comparable with Brandl et al. (2020) for spruce, beech and Scots pine; oak species were not separated in this study. This study found higher long-term summer temperature related with higher mortality for pedunculate oak, whereas in the case of sessile oak, higher continentality led to reduced mortality. The ability of pedunculate oak to withstand continental conditions was well reflected in its eastern distribution range far into Russia, but the distribution of sessile oak did not extend as far, so the result was not evident. Oak decline has been reported to be associated with human, environmental and biotic factors, such as lowering groundwater table, absence of flooding, pollution, non-adapted silvicultural practices and climate change (Eaton et al. 2016). Furthermore, pedunculate oak also grows on heavier soils, in wet lowlands and damp areas by streams and rivers, tolerating periodic flooding. Thus, vitality of this species might be more dependent on soil conditions and water table (e.g. Kostić et al. 2021). The overall pattern of relation between sessile oak and climate variables differed from the rest of the species set which might be due to decreasing mortality trends in the decade studied.

Long-term averages for precipitation either increased (pedunculate oak, spruce, Scots and black pine) or decreased (hornbeam, Austrian and sessile oak) odd ratios for mortality. We can only speculate why higher summer precipitation is related to higher mortality in case of pedunculate oak, a drought tolerant species (Eaton et al. 2016; Niinemets and Valladares 2006). A common problem for oaks is a combination of mildew of the second leaf flushing after massive defoliation (Eaton et al. 2016). The effect for spruce is very weak, which is in line with Brandl et al. (2020) who found no precipitation effect for this species. Black pine provenances might suffer from fungal diseases (winter temperature and summer rain were both positively correlated with *Diplodia* colonization on cones in France, and become virulent after drought periods (Fabre and Piou 2011)). These findings are supported by results of this study. Higher spring precipitation showed odds ratio above 2 for Scots pine. This again might be linked to *Diplodia* or other fungi or to wet snow (most likely in late autumn and early spring) and subsequent crown breakage that starts a decline spiral (Nykänen et al. 1997).

Besides long-term climate data (predisposing factors) that are related to diffuse background mortality, climate anomalies (pulse events) showed important effects on mortality and can be seen as inciting factors (Manion 1991; Manion and Lachance 1992; Sturrock et al. 2011; Senf et al. 2020). Anomalies are known to reduce growth (Thurm et al. 2016; Vanoni et al. 2016) or vitality measured for example as annual defoliation (Eickenscheidt et al. 2019). Mortality occurs when physiological thresholds are exceeded (Schuldt et al. 2020; Senf et al. 2020; Vanoni et al. 2016), which is likely under extremes as “plants have limited physiological potential to respond to rapid changes in the environment” (Choat et al. 2018). Additionally, both climate and weather promote systemic effects linked to vitality decline, such as pests or diseases that favor warmer temperatures (Sturrock et al. 2011). The impact of weather anomalies on forest insects is complex, but anomalies such as unusually warm or dry conditions can serve as initial eliciting factors for pest outbreaks (Raffa et al. 2008).

With the exception of European hornbeam, two to three year averages of temperature anomalies for the warmest month, summer, spring and vegetation periods were most influential in the models, indicating lag effects and longer lasting stress and/or favorable weather conditions for insects or fungi. Summer, vegetation period or annual precipitation sum anomalies were significantly selected as two to three year averages with the exception of spruce. In contrast, the effects of winter anomalies (temperature and precipitation) had shorter time spans and therefore often a more direct relation with mortality of the subsequent assessment which takes place in summer. This might indicate different processes: stress via heat and drought which starts a decline and a direct effect on insects or pathogens but also change in phenology that increases risk for late frost damage or insect damage. A review on phenological form of pedunculate oak states that there are two phenological types – early and late flushing – and that the late form of oak is more resistant to spring frosts and insect damage (Utkina and Rubtsov 2017).

This study found that warmer than average winters reduced mortality for birch, Austrian and sessile oak, whereas warmer coldest month temperatures increased risk for pedunculate oak, spruce, Scots pine (both weak effects), beech and black pine. Neumann et al. (2017) found that a warmer than average maximum temperature of winter season increased mortality whereas warmer than average minimum temperature of winter season decreased mortality in the 2000-2012 period in Europe, which was interpreted that there is a reduction in cold-induced mortality. Warmer winters, earlier flushing, higher risk of late frost and higher survival of insects (Allen et al. 2010), and reduced cold-induced mortality under harsh environments might be effects related to findings of this study. Still, it is not clear why birch as an extremely cold hard species is harmed by harsher winters. Austrian and sessile oak show a lower cold hardiness and therefore might benefit (hardiness ranking according to Roloff and Grundmann (2008)). Dietze and Moorcroft (2011) found differing response for minimum winter temperature for differing plant functional types in North America, which highlights the regional and species-specific differences between effect and response that was also found here. Northern types showed higher mortality under colder winter (ice damage or freeze embolism), a result found in this study for birch, Austrian and sessile oak, the latter two not growing in Northern Europe.

In line with several studies (Brandl et al. 2020; Maringer et al. 2021; Allen et al. 2010, 2015; Ciais et al. 2005) and expectations of this study, warmer temperatures in the previous year increased mortality risk dramatically for five out of nine species. This was strongest for species that dominate Northern European forests (birch, spruce, Scots pine) either because they are not adapted to this situation or the link to insects and pests, which benefit from reduced temperature limitation. The overall effect with the coupling of stress and population dynamics driven by drought and heat is well documented in case of bark beetle infesting spruce (Anderedd et al. 2015; Marini et al. 2018; Raffa et al. 2008). This study found a contrasting result for sessile oak, beech and black pine (higher temperatures in spring or warmest month related to reduced mortality). This effect should be expected in regions where warming reduces stress, for example, Jolly et al. (2005) described how a longer vegetation period resulted in better tree growth in the Swiss Alps. Black pine growth improves with positive spring water balance (Marqués et al. 2016), so positive spring anomalies might reduce mortality when accompanied with precipitation or when growth limits due to low temperatures are reduced. Findings for spruce, beech but not for Scots pine are in line with results of Nothdurft (2013), though the later study focused on only one region in Germany, and was based on a small data set and a different period.

Effects of precipitation anomalies split up between winter and non-winter periods (vegetation period, summer or annual precipitation). Three out of four species (hornbeam, pedunculate oak, black pine) suffered from higher winter precipitation, a result that normally is related to wind throw risk due to water saturated soils when it coincides with high winds (reviewed by Mitchell 2013), which was excluded from the data set of this study. Annual, summer or vegetation period precipitation anomalies had to be taken as variables of the previous years, as crown condition survey takes place in summer. Therefore, results do not reflect only direct drought effects, but also lagged consequences, which was apparent from the steep increase of mortality following 2018 for spruce and birch in data of this study (see also Brun et al. 2020; Sent et al. 2020). Neumann et al. (2017) pointed out that variability between summer and spring anomalies of the previous period are drivers of mortality (wet summer and dry spring). A reason for this might be the timing of bud formation in mid-summer, so that the number of leaves is higher after a wet period which increases water demand in a subsequent drought in the following spring or summer (Meier and Leischner 2008). Only Austrian oak witnessed a clear beneficial effect of higher precipitation, which is in accordance with drought susceptibility of this species (Colangelo et al. 2018). Effects of interaction between temperature and precipitation variables for spruce and Scots pine could be interpreted in the way that rising temperatures in the vegetation period can be compensated by higher precipitation.

Interpretation of observed results regarding the impact of aspect on mortality have limitations. Aspect and slope were not significant in other studies, e.g. Maringer et al. (2021). The data on aspect did not incorporate information on slope, however it is understood that steep slopes in southern directions are warmer (Reger et al. 2011). Effects of aspect could be hypothesized to be that southwest slopes are drier and warmer and north-facing slopes are colder and even prone to frost-drought (Tranquillini 1982). In the models, beech and oaks tended to have lower mortality on south and west facing sites, whereas spruce exhibited this pattern only for western slopes. For black pine, the highest mortality estimates were calculated for sites facing east and west, while it was northeast and northwest for Scots pine. These are of course just generalized results of the models under the caveat that each species will favor different aspects, depending on altitude and latitude.

Neumann et al. (2017) found that warm summers as well as high seasonal variability in precipitation increased the likelihood of tree death, and that age was an important driver of mortality for European tree species. Brandl et al. (2020) found increasing mortality risk with age for spruce and beech, but not so clear dependencies for oaks and Scots pine. This study found decreasing mortality with age for pine and oak species, and a similar, but weak trend for spruce. For beech, risk first diminished and then increased with age. For the short-lived pioneer species silver birch as well as short-lived hornbeam, age increased mortality considerably (Leuschner and Ellenberg 2017). Trees in uneven aged stands had a higher mortality compared to the reference group 41-60 years for Norway spruce, European beech, and silver birch, but lower mortality for Scots pine and sessile and pedunculate oak. Age effects might be related with development stages of stands and age-effects were leveled out between species when survival probabilities were displayed over tree height in the study by Maringer et al. (2021).

The effects of extreme and long-lasting droughts and heat waves on mortality according to the second hypothesis is related with the short-term effects of annual anomalies. Europe has experienced a series of extremely hot and dry summers (2003, 2010, 2013, 2015, 2018) (Hanel et al. 2018; Buras et al. 2020). Droughts were driven by heatwaves and precipitation deficits in the vegetation period, affected certain parts of Europe and the effects on vegetation was modulated by the weather conditions in the preceding winter or subsiding seasons. For example, the 2015 summer was the hottest since 1950 across a large part of eastern and south-western Europe, but in terms of annual precipitation deficit “the severity of the 2015 drought was possibly limited due to the wet preceding winter over large part of Europe” (Hanel et al. 2018). Buras et al. (2020) described 2018 as characterized by a climatic dipole, “featuring extremely hot and dry weather conditions north of the Alps but comparably cool and moist conditions across large parts of the Mediterranean”. These spatio-temporal patterns have consequences on mortality as species are not evenly distributed and affected in single events. This was reflected in data here that exhibited a steep increase in annual mortality after 2018 for spruce, birch, and also beech, but showed no signs of increased mortality for species with a Mediterranean distribution like Austrian oak, and black pine or species that encompass at least considerable distributions in the Mediterranean like sessile oak. The effects of the drought 2018 on spruce and beech in Germany were enhanced by consecutive drought years and led to increased mortality rates (Obladen et al. 2021). This aggravating effect of consecutive unfavourable weather conditions with an increase in soil moisture drought (Hanel et al. 2018) explains why there was no drop in annual mortality for Norway spruce or birch after 2018. Strong drought-legacy effects in 2019, with missing physiological recovery, that leave trees highly vulnerable to secondary insect or fungal pathogen attacks were also reported in (Schuldt et al. 2020; Moravec et al. 2021). In contrast, there was no effect of the hot and dry summer of 2013 (Hanel et al. 2018) visible in data of this study. Apart from Austrian oak there was no increased annual mortality in 2013 or 2014. Thus, we can conclude that short-term effects of annual anomalies influence tree mortality but the severity of the effects is also shaped by previous (Hanel et al. 2018) and subsequent (Schuldt et al. 2020) weather conditions.

The third hypothesis is difficult to judge as it is somewhat contradicting to the assumption of an individual response of species to environmental stress. Nevertheless, this hypothesis was formulated due to spatial aspects in species distribution. Figure 2 can be interpreted for grouping of species according to effects. It revealed similarities between some broadleaved species of warmer climates and lowlands (Austrian and sessile oaks), and species that also occur in higher elevations and colder climates (spruce, pine, and pedunculated oak) with distinct cold hardiness, as described by Kreyling et al. (2012) for *P. nigra. Q. cerris* forms mixed stands together with *Q. petraea* in Austria and southern Slovakia as outposts in the Pannonian region (Leuschner and Ellenberg 2017). Beech and hornbeam appear to fall between those two groups. The pioneer species silver birch with only three significant climate or weather variables impacting its mortality is hard to classify with other species. Species tolerating warmer conditions possess mechanisms to withstand heat and drought to a certain degree. Oak seedlings, for example, exposed to drought have been shown to have adapted their growth and xylem structure to improve drought resistance, for example, via reduction of latewood vessel size (Vander Mijnsbrugge et al. 2020). Grouping according to distribution was also performed for North American species into plant functional traits with subsequent mortality patterns analyzed by Dietze and Moorcroft (2011), who distinguished between angiosperms and gymnosperms. Shade tolerance and abundant species groups were further separated into Northern and Southern ranges according to established differences in climatic tolerances. Only the distributional aspect of this classification might be appropriate for this study.

The low-land and higher elevation groups differed clearly in the effects of winter temperature anomalies. Broadleaved species of warmer climates might benefit from a lower risk of severe frost damage, a longer vegetation period with an earlier flushing to allow for use of soil water stocks from winter precipitation (e.g. in Mediterranean eco-systems). The negative effect of warmer (and wetter) winters for conifers is often related to wind throw (Maringer et al. 2021), an effect we excluded from our data set. Still, trees can be injured by snow break and there is a feedback on pathogens and pests with winter anomalies. Besides that, warmer winters are known to increase the likelihood for late spring frost damage in frost prone regions as trees begin wood formation, leaf release and flowering weeks earlier compared to the mid-twentieth century (Puchałka et al. 2016, 2017). The greater frequency of late spring frosts is related to climate warming (Sangüesa-Barreda et al. 2021).

A limitation of the study was the low event rate of mortality that affects the power for complex multivariable statistical models to detect weather and climate impacts, particularly concerning interactions among risk factors. These limitations are common to any study of rare events. Mortality is assessed annually, necessitating the use of logistic regression methods for interval censored data as described in the methods. While exact mortality dates are available for humans and animals, and allow more accurate modeling, such as by the Cox proportional regression model, they do not exist for trees, where annual assessment might be considered the least coarse option available for mortality assessments. The logistic regression methods outlined in this study for binary outcomes are easily transportable to other forest monitoring scenarios, they produce interpretable odds ratios providing quantitative effect sizes in addition to statistical significance.

## 4. Materials and Methods

Tree mortality data across Europe for the decade spanning 2011-2020 were provided by the International Co-operative Programme on Assessment and Monitoring of Air Pollution Effects on Forests (ICP Forests). Level I data on systematic 16×16 km grids were pooled with Level II intensive monitoring plots (Michel et al. 2018). Although different plot designs were used by these monitoring levels, they shared the same method of assessment, and data were pooled to tree-year observations. Individual tree crown condition has been assessed annually by the ICP Forests since the 1980’s, depending on the country. Since 2011 Level I trees were reported with an additional removal or mortality code, enabling distinction between trees that died and trees that were removed through management operations. Even though this removal code was established for Level II plots earlier, tree observations starting from 2011 were used for both data sets to have a consistent time range. The mortality events were classified as those due to biotic and abiotic causes, as well as standing dead trees where the cause of death was unknown (Table S10). Planned utilization (forest management) and fallen trees were excluded. The status of fallen trees is in many cases unknown due to an ambiguity in the ICP Forests coding system. Overall 746,478 tree-year observations from 32 countries were analyzed. These observations came from 130,018 trees from 8618 plots over the ten-year period studied.

For all plots, the plot information aspect (Flat, South, West, …) and stand mean age were used. Since individual tree age was not recorded, mean stand age was used as proxy, provided in 20-year classes, with one category for irregular stands. The class of irregular mean stand age refers to uneven-aged stands containing two or more distinct age classes. For inclusion analyses, only codominant and dominant trees older than 40 years were considered in order to focus on climatic drought effects and neglect competition effects. Only the most common European tree species were included for analysis. Specifically, only the nine species with more than 10,000 tree-year observations and at least 100 dead trees in the sample were included for analysis, shown in Table 1.

Tree mortality data were combined with weather data at plot location from ClimatEU (Marchi et al. 2020), which mapped climate indices at a 2.5 arcmin grid (approximately 5 km) over Europe. An advantage was that the weather data was adjusted for the provided elevation by the data software applying partial differential equations. Bioclimatic variables such as temperature, precipitation and continentality were considered for different time periods. A complete list of all variables can be found in Table 2. Annual and seasonal weather characteristics of the year of reported tree death and the three previous years were used. Winter was defined as December of the previous year to February, spring as March to May, summer as June to August and the vegetation period as May to September. To account for the general environment a tree grew in, anomalies of the weather variables were calculated at plot level by subtracting the 30-year mean of the years 1981 to 2010 of the seasonal value. The climate normal of this 30-year period was selected because it was experienced by all tree stands. Annual, summer and vegetation period variables from the year before tree death were used as risk factors, since crown condition assessments were performed in summer. As further predictors, the cumulative means of the previous three years were calculated, starting from the first concerned year depending on summer or winter variables. Climate normals were used as predictors for the general environment, with additionally the difference of mean warmest and mean coldest temperature, resulting in 50 weather characteristics of interest.

To investigate yearly trends and corresponding weather effects, mortality was reported as the number of observed dead trees in one year divided by all observed trees for that year and compared with the annual weather variables averaged for all available trees for one year. All annual measures were standardized to mean 0 and standard deviation 1 over the 10-year period. Mortality was shown for all tree species together and combined with the weather trends described in an exploratory fashion. Associations in annual trends does not necessary imply significant associations in the aggregated logistic regression models, which pool all annual observations and adjust for all potential confounding variables, such as age. Individual annual trend analyses do not have sufficient sample sizes to perform adjustment for confounding variables, and would furthermore lead to high amounts of multiple comparisons and spurious statistical associations. Pooling annual data into a single logistic regression for investigating the multiple conflicting effects on tree mortality obtains sufficient sample size and power for detecting significant effects of weather and climate factors regardless of the year(s) in which they occurred.

Because tree death is only observed yearly, meaning that the specific date of the death is not observed and only the cumulative survival over the year, ordinary survival models that do not account for interval censoring, such as the Cox proportional hazards model, are not appropriate. Abbott (1985) showed that logistic regression can be used to approximate interval censored survival data analysis and following Boeck et al. (2014), this approach was adopted for the analysis of yearly tree mortality in this paper. Age and aspect were only considered as predictor in the final model for a tree species, if tree-year observations were observed for all categories. The interpretation of the coefficients from logistic regression is as follows. The exponential of the coefficient for a particular covariate in the model is the odds ratio (OR) for experiencing a tree mortality for a unit increase in the covariate according to its unit of measurement, for example for an increase in one degree Celsius (°C) temperature or its anomaly. For categorical variables, the interpretation is the change in OR for each category compared to a reference category. As unit increase we chose one °C for temperature and 50 mm for precipitation variables, as reasonable difference over the large gradient of Europe. For model evaluation we calculated the area under the receiver operating characteristic curve (AUC) as accuracy measure, were an AUC greater 50% indicates ability to detect tree mortalities (Mandrekar 2010).

Risk factor significance was assessed via likelihood ratio tests, with all tests performed at the 0.05 level of significance. For selecting the optimal model, forward selection starting with an intercept only model was performed as follows. Conditioned on the current model, addition of a variable with the smallest p-value among all variables not in the model was considered, and included if the p-value fell below 0.05. Multi-collinearity among risk factors was reduced by building variable groups according to similarity following approaches in species distribution modelling by Mellert et al. (2011) and Thurm et al. (2018). Therefore, variable groups were formed as shown in Table 2. For each group of qualitatively similar variables, such as multiple measures of precipitation, only the best single variable significantly improving the model in terms of lowest p-value was included in the model. The disadvantage of incorporating both long-term average (climate normal) and immediate anomaly (weather data) climate signals in models is their correlation, which leads to interference within the regression models and sometimes, contradictory effects. Interaction terms were included in the models when statistically significant in order to gain insight into the interfering effects of the correlated risk factors. This procedure led to models with a maximum of five main effects from climate or weather variables plus potential interaction terms among them. For each species the final model was evaluated at each location in the data set with the corresponding weather factors and visualized via prediction maps alongside locations of observed dead trees. All computations and figures were made with the statistical package R version 4.1.1 (R Core Team 2021).

## 5. Conclusions

Different types of mortality – background or disturbances - are related to different variables and time scales. Multi-year stress events do hit managed forests with tremendous economic loss and do hamper fulfillment of ecosystem services as e.g. carbon storage. Tree mortality models and predictions from this study, combined with climate change projections, could be used to improve forest management planning. It is important to stress that climate warming increases the risk of mortality of all studied tree species with less severity in case of sessile oak, and this understanding should be the baseline of future forest management plans throughout Europe. Optimal tree species may be selected for planting dependent upon regional climates. Still, it is important to understand differences related to mortality between managed or hugely human influenced forests and natural forests. The individual tree logistic regression models should be updated annually with new mortality data, including precise age estimates, which would be especially beneficial for increasing the statistical power to detect effects for other tree species, as well as incorporate recent weather events, thus maximally preparing for an uncertain future under climate change. Remote sensing techniques add additional value for future data on individual tree mortality.

## Supplementary Materials

Annual values for weather variables and mortality are shown in Figure S1 and S2. Model estimates and evaluations for all species are presented in Tables S1-S9 and Figures S3-S10. Mortality classification is shown in Table S10.

## Declarations

### Availability of data and material

Restrictions apply to the availability of study data. Forest data was obtained from ICP Forests and are available at http://icp-forests.net/ with the permission of ICP Forests. Weather data is publicly available as described in Marchi et al. (2020).

### Code availability

Code can be provided by the corresponding author on request.

### Author Contributions

Formal analysis: Matthias Neumair; Methodology: Matthias Neumair, Donna Ankerst and Wolfgang Falk; Project administration: Donna Ankerst and Wolfgang Falk; Supervision: Donna Ankerst and Wolfgang Falk; Visualization: Matthias Neumair; Writing – original draft: Matthias Neumair, Donna Ankerst and Wolfgang Falk; Writing – review & editing: Nenad Potočić, Volkmar Timmermann, Mladen Ognjenović and Susanne Brandl.

## Acknowledgments

Not applicable.

