## supplementary for "Drought effects of annual and long-term temperature and precipitation on mortality risk for 9 common European tree species"

**Supplementary Materials**

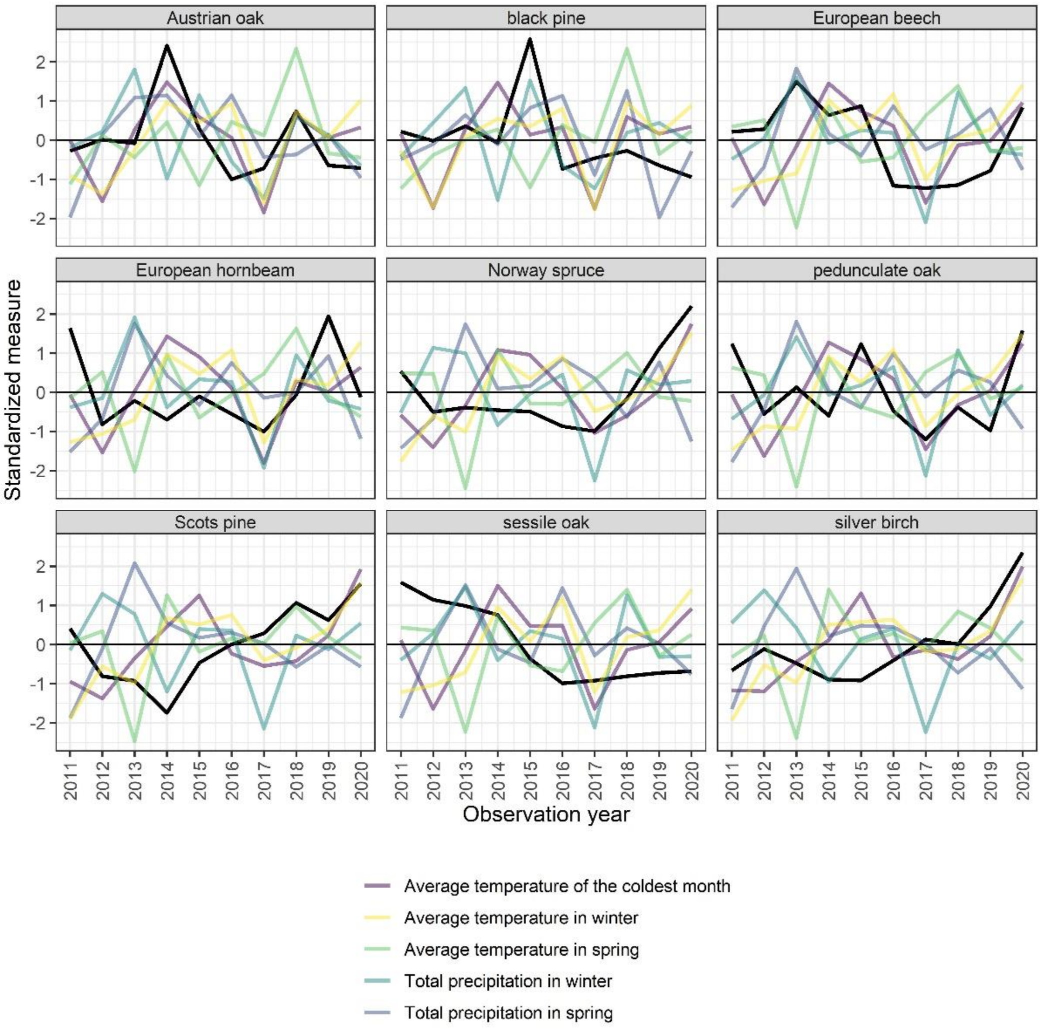


**Fig S1** Annual values in winter and spring weather variables and mortality (black line) for the individual species. All variables have been standardized by subtraction of their means and divided by their standard deviations

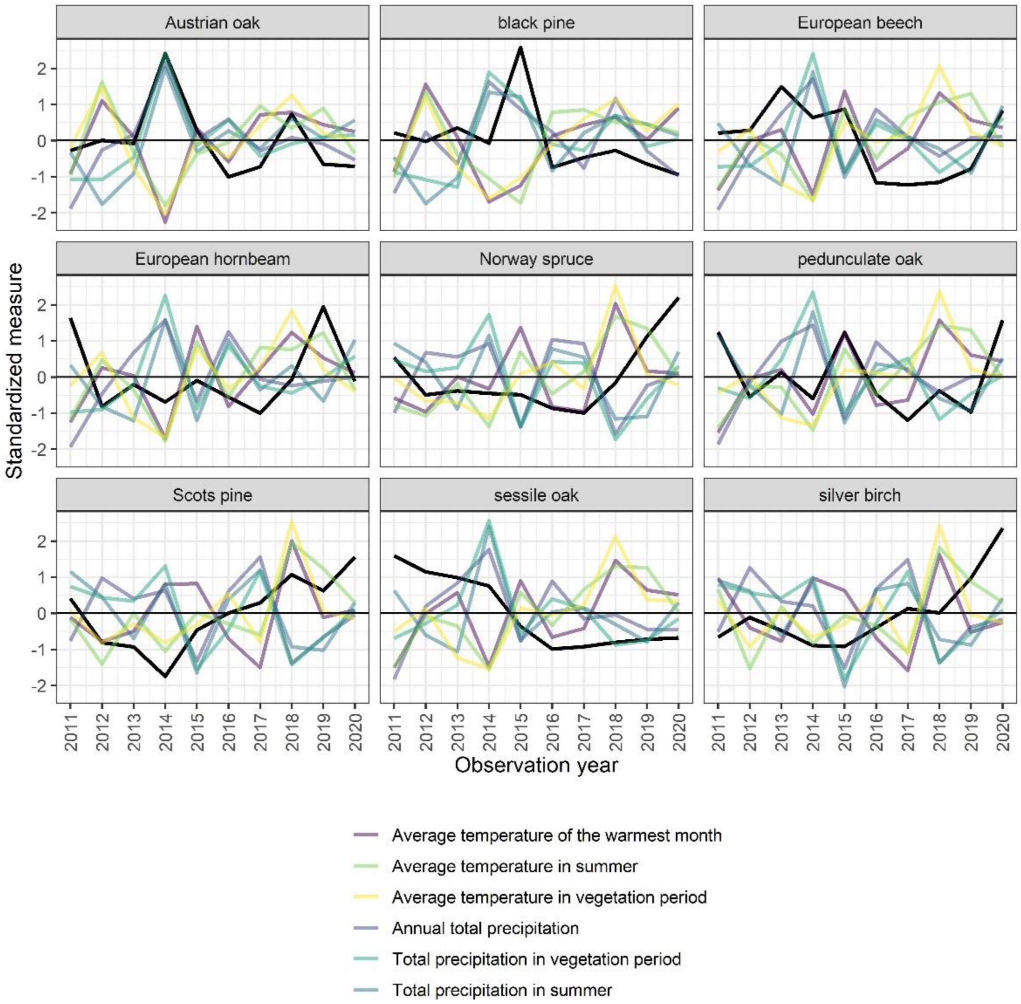


**Fig S2** Annual values in summer weather variables and mortality (black line) for the individual species. All variables have been standardized by subtraction of their means and divided by their standard deviations

European beech

**Table S1.** Logistic regression model estimates for European beech.

| Variable | Odds ratio | Confidence interval | P-value |
|---|---|---|---|
| Mean stand age (Reference 41 -60) | | | 0.003 |
| 61 – 80 | 1.03 | 0.74 – 1.45 | |
| 81 – 100 | 0.85 | 0.60 – 1.21 | |
| 101 - 120 | 0.91 | 0.63 – 1.33 | |
| >120 | 1.53 | 1.10 – 2.14 | |
| Irregular stands | 1.37 | 0.88 – 2.11 | |
| Aspect (Reference Flat) | | | < 0.0001 |
| North | 1.31 | 0.89 – 1.98 | |
| North-east | 0.71 | 0.45 – 1.14 | |
| East | 1.58 | 1.04 – 2.42 | |
| South-east | 0.56 | 0.30 – 1.02 | |
| South | 0.78 | 0.47 – 1.27 | |
| South-west | 0.52 | 0.28 – 0.93 | |

| | | | |
|---|---|---|---|
| West | 0.85 | 0.54 – 1.33 | |
| North-west | 1.42 | 0.94 – 2.19 | |
| Temperature difference between average temperature of the coldest and the warmest month (continentality) 30-years average (°C) | 1.22 | 1.15 – 1.29 | < 0.0001 |
| Average temperature of the coldest month anomaly at reported tree mortality year (°C) | 1.19 | 1.13 – 1.26 | < 0.0001 |
| Total precipitation in winter anomaly average of the reported tree mortality year (50mm) | 0.74 | 0.56 – 0.96 | 0.02 |
| Average temperature of the warmest month anomaly average of 3 previous years before reported tree mortality year (°C) | 0.50 | 0.37 – 0.67 | < 0.0001 |
| Interaction between total precipitation in winter anomaly average of the reported tree mortality year and average temperature of the warmest month anomaly average of 3 previous years before reported tree mortality year (°C * 50mm) | 1.39 | 1.15 – 1.66 | 0.0005 |

Black pine

**Table S2.** Logistic regression model estimates for Black pine. Trees older than 120 years were excluded due to lack of tree mortalities in this group.

| **Variable** | **Odds ratio** | **95% Confidence interval** | **P-value** |
|---|---|---|---|
| Mean stand age (Reference 41 -60) | | | < 0.0001 |
| 61 – 80 | 0.43 | 0.24 – 0.75 | |
| 81 – 100 | 0.33 | 0.15 – 0.63 | |
| 101 - 120 | 0.33 | 0.16 – 0.65 | |
| Irregular stands | 0.45 | 0.22 – 0.86 | |
| Aspect (Reference Flat) | | | 0.003 |
| North | 2.36 | 0.83 – 8.46 | |
| North-east | 2.96 | 1.07 – 10.51 | |
| East | 4.42 | 1.69 – 15.13 | |
| South-east | 3.95 | 1.27 – 14.70 | |
| South | 4.92 | 1.83 – 17.12 | |
| South-west | 1.64 | 0.55 – 6.00 | |
| West | 1.95 | 0.58 – 7.55 | |
| North-west | 2.26 | 0.76 – 8.19 | |
| Average temperature in summer 30-years average (°C) | 1.38 | 1.22 – 1.55 | < 0.0001 |
| Total precipitation in summer 30-years average (50mm) | 1.46 | 1.34 – 1.59 | < 0.0001 |
| Average temperature of the coldest month anomaly average of the reported tree mortality and 1 previous years (°C) | 1.38 | 1.19 – 1.61 | < 0.0001 |
| Average temperature in spring anomaly average of the reported tree mortality year and 2 previous years (°C) | 0.47 | 0.30 – 0.73 | 0.0007 |
| Total precipitation in winter anomaly of the reported tree mortality year (50mm) | 1.16 | 1.04 – 1.29 | 0.008 |

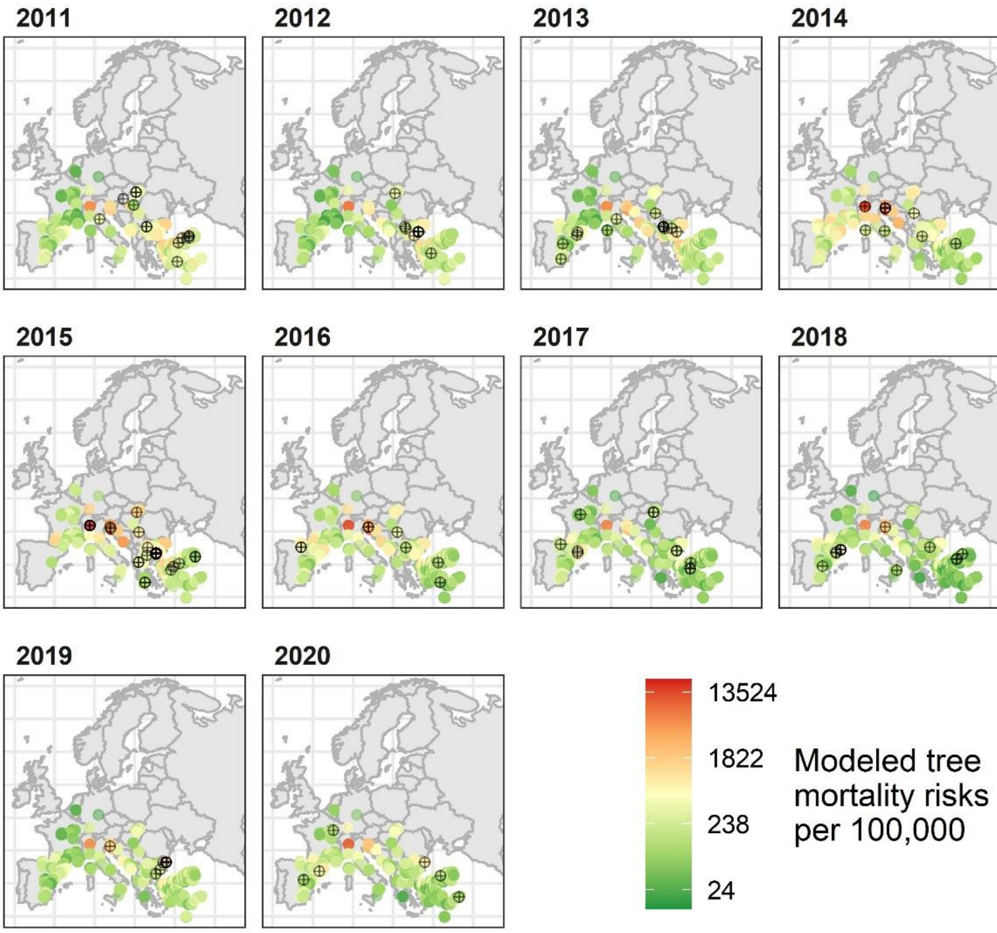


**Fig S3** Spatial evaluation of modelled tree mortality for Black pine for all analyzed ICP Forests plots on a logarithmic color scale. Black points indicate observations of dead trees (n=132)

Norway spruce

**Table S3.** Logistic regression model estimates for Norway spruce.

| Variable | Odds ratio | 95% Confidence interval | P-value |
|---|---|---|---|
| Mean stand age (Reference 41 -60) | | | < 0.0001 |
| 61 – 80 | 0.67 | 0.55 – 0.80 | |
| 81 – 100 | 0.90 | 0.76 – 1.08 | |
| 101 - 120 | 1.06 | 0.88 – 1.27 | |
| >120 | 0.79 | 0.64 – 0.96 | |
| Irregular stands | 1.19 | 0.89 – 1.56 | |
| Aspect (Reference Flat) | | | < 0.0001 |
| North | 1.47 | 1.22 – 1.76 | |
| North-east | 1.31 | 1.05 – 1.63 | |
| East | 1.17 | 0.94 – 1.45 | |
| South-east | 1.47 | 1.19 – 1.80 | |
| South | 1.71 | 1.41 – 2.06 | |
| South-west | 0.73 | 0.56 – 0 95 | |
| West | 0.68 | 0.53 – 0.87 | |

| | | | |
|---|---|---|---|
| North-west | 0.55 | 0.42 – 0.72 | |
| Total precipitation in vegetation period 30-years average (50mm) | 1.06 | 1.04 – 1.08 | < 0.0001 |
| Temperature difference between average temperature of the coldest and the warmest month (continentality) 30-years average (°C) | 1.10 | 1.07 – 1.13 | < 0.0001 |
| Average temperature of the coldest month anomaly at reported tree mortality year (°C) | 1.04 | 1.01 – 1.06 | 0.004 |
| Average temperature in vegetation period anomaly average of 2 previous years before reported tree mortality (°C) | 2.28 | 2.01 – 2.60 | < 0.0001 |
| Total precipitation in vegetation period anomaly of the previous year before reported tree mortality (50mm) | 1.27 | 1.22 – 1.33 | < 0.0001 |
| Interaction between average temperature in vegetation period anomaly average of 2 previous years before reported tree mortality and total precipitation in vegetation period anomaly of the previous year before reported tree mortality (°C * 50mm) | 0.81 | 0.77 – 0.85 | < 0.0001 |

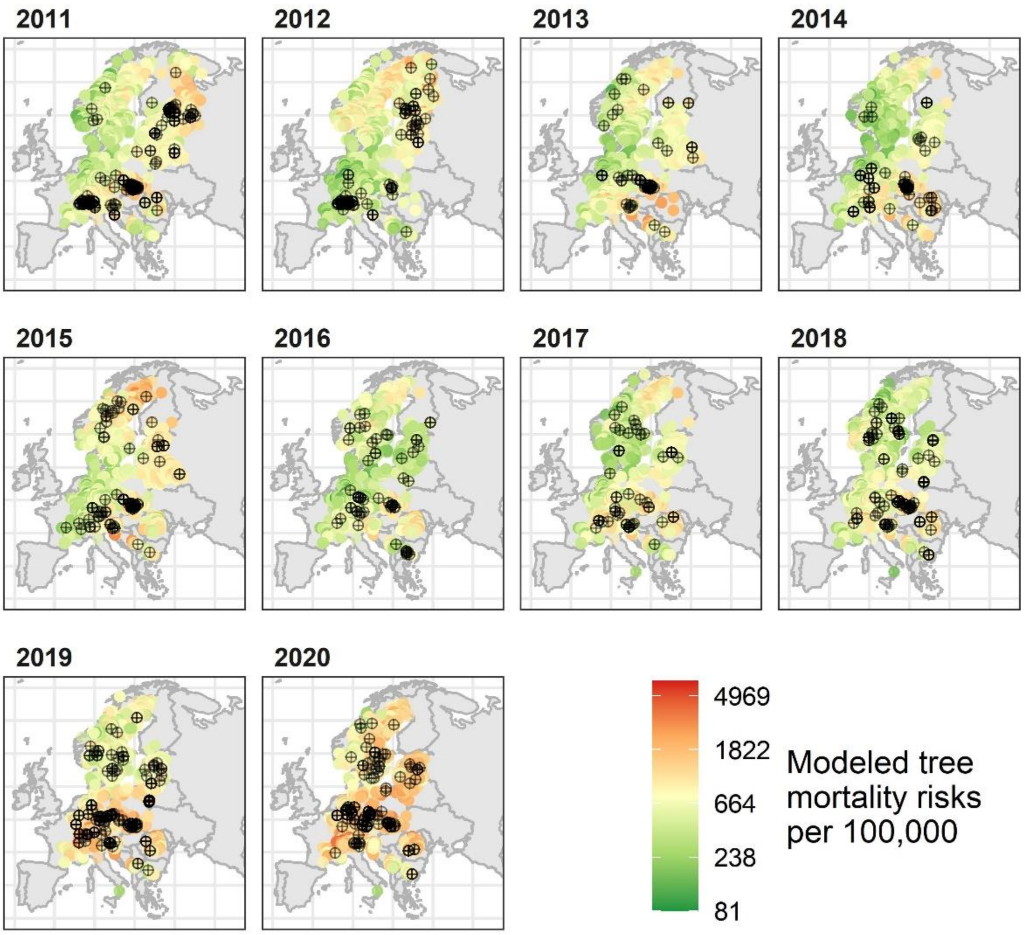


**Fig S4** Spatial evaluation of modelled tree mortality for Norway spruce for all analyzed ICP Forests plots on a logarithmic color scale. Black points indicate observations of dead trees (n=1399)

Scots pine

**Table S4.** Logistic regression model estimates for Scots pine.

| Variable | Odds ratio | 95% Confidence interval | P-value |
|---|---|---|---|
| Mean stand age (Reference 41 -60) | | | < 0.0001 |
| 61 – 80 | 0.58 | 0.49 – 0.69 | |
| 81 – 100 | 0.64 | 0.53 – 0.77 | |
| 101 - 120 | 0.56 | 0.43 – 0.72 | |
| >120 | 0.73 | 0.55 – 0.96 | |
| Irregular stands | 0.81 | 0.55 – 1.15 | |
| Aspect (Reference Flat) | | | < 0.0001 |
| North | 1.29 | 0.95 – 1.72 | |
| North-east | 6.93 | 5.61 – 8.54 | |
| East | 2.10 | 1.44 – 2.97 | |
| South-east | 1.13 | 0.71 – 1.71 | |
| South | 1.56 | 1.18 – 2.03 | |
| South-west | 1.21 | 0.83 – 1.69 | |
| West | 2.76 | 2.11 – 3.55 | |
| North-west | 2.73 | 2.10 – 3.52 | |
| Average temperature of the warmest month 30-years average (°C) | 1.28 | 1.15 – 1.43 | < 0.0001 |
| Total precipitation in spring 30-years average (50mm) | 2.10 | 1.27 – 3.42 | 0.004 |
| Interaction between average temperature of the warmest month 30-years average and total precipitation in spring 30-years average (°C * 50mm) | 0.95 | 0.92 – 0.98 | 0.0004 |
| Average temperature in vegetation period anomaly average of 2 previous years to the reported tree mortality year (°C) | 2.89 | 2.45 – 3.42 | < 0.0001 |
| Annual total precipitation anomaly average of 3 previous years to the reported tree mortality year (50mm) | 1.79 | 1.57 – 2.05 | < 0.0001 |
| Average temperature of the coldest month anomaly average of the reported tree mortality and 3 previous years (°C) | 1.03 | 0.96 – 1.11 | 0.4 |
| Interaction between average temperature in vegetation period anomaly average of 2 previous years to the reported tree mortality year and annual total precipitation anomaly average of 3 previous years to the reported tree mortality year (°C * 50mm) | 0.80 | 0.70 – 0.91 | 0.0008 |
| Interaction between average temperature of the coldest month anomaly average of the reported tree mortality and 3 previous years and annual total precipitation anomaly average of 3 previous years to the reported tree mortality year (°C * 50mm) | 1.17 | 1.11 – 1.24 | < 0.0001 |

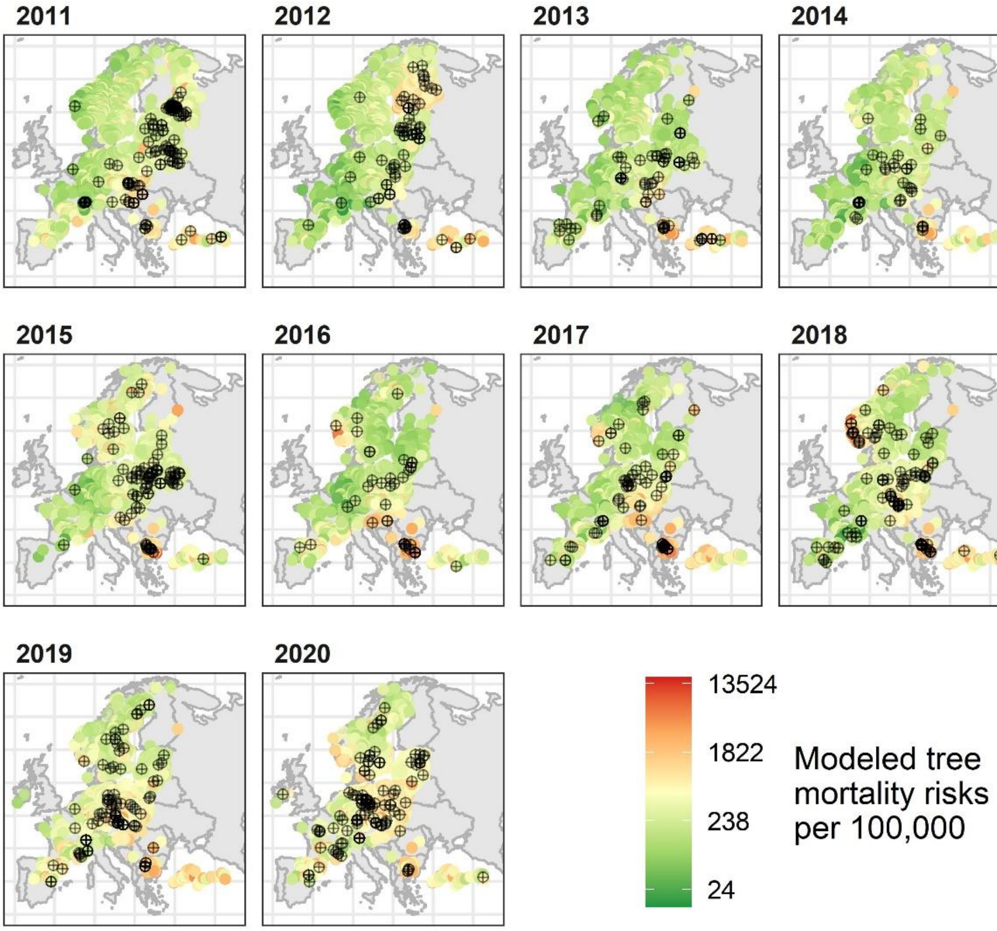


**Fig S5** Spatial evaluation of modelled tree mortality for Scots pine for all analyzed ICP Forests plots on a logarithmic color scale. Black points indicate observations of dead trees (n=953)

Silver birch

**Table S5.** Logistic regression model estimates for Silver birch.

| Variable | Odds ratio | 95% Confidence interval | P-value |
|---|---|---|---|
| Mean stand age (Reference 41 -60) | | | 0.0004 |
| 61 – 80 | 1.77 | 1.16 – 2.74 | |
| 81 – 100 | 1.87 | 1.01 – 3.33 | |
| 101 - 120 | 3.53 | 1.58 – 7.12 | |
| >120 | 4.74 | 2.25 – 9.27 | |
| Irregular stands | 1.68 | 0 82 – 3.36 | |
| Average temperature in winter 30-years average (°C) | 1.15 | 1.08 – 1.23 | < 0.0001 |
| Average temperature in vegetation period anomaly average of 2 previous years before reported tree mortality (°C) | 5.44 | 3.38 – 8.91 | < 0.0001 |
| Average temperature in winter anomaly at reported tree mortality year (°C) | 0.90 | 0.82 – 0.99 | 0.03 |

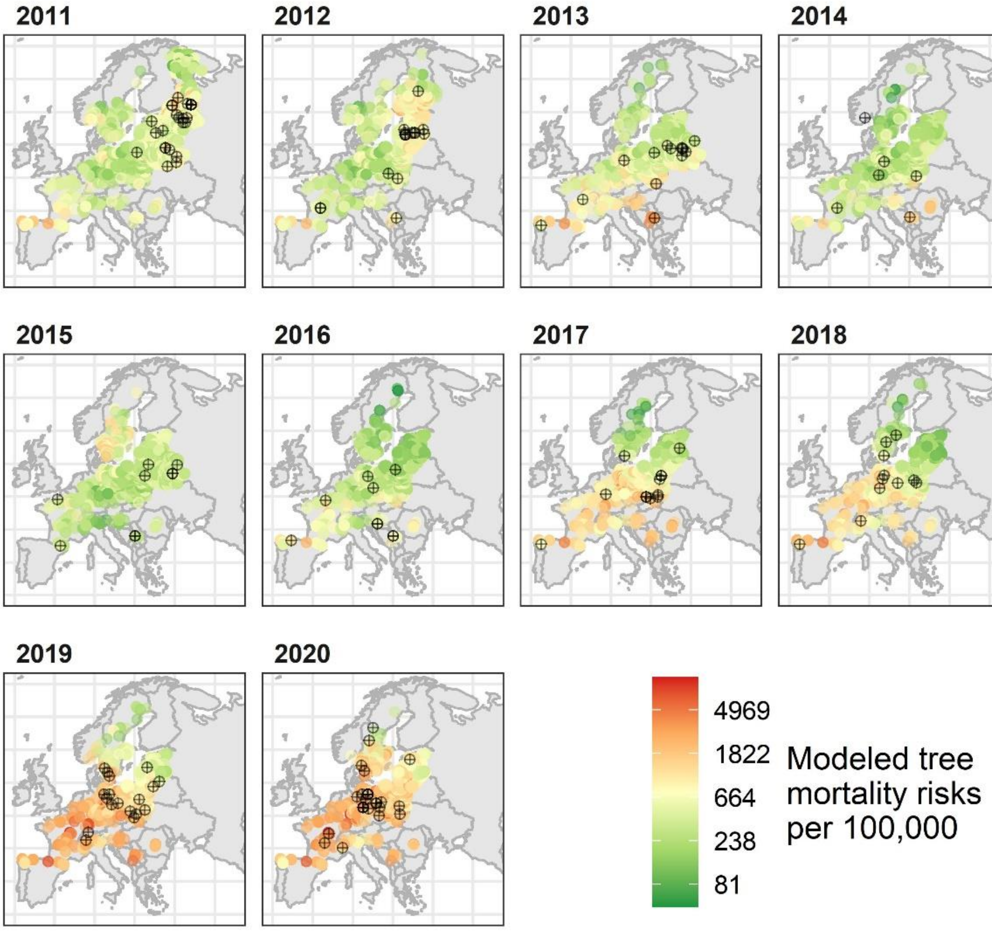


**Fig S6** Spatial evaluation of modelled tree mortality for silver birch for all analyzed ICP Forests plots on a logarithmic color scale. Black points indicate observations of dead trees (n=145)

Austrian oak

**Table S6.** Logistic regression model estimates for Austrian oak.

| Variable | Odds ratio | 95% Confidence interval | P-value |
|---|---|---|---|
| Mean temperature of spring 30-years average (°C) | 1.60 | 1.35 – 1.92 | < 0.0001 |
| Precipitation sum of winter 30-years average (50mm) | 0.85 | 0.72 – 0.98 | 0.02 |
| Average temperature in winter anomaly average of the reported tree mortality year and 3 previous years(°C) | 0.60 | 0.42 – 0.86 | 0.005 |
| Total precipitation in summer anomaly average of 3 previous years to the reported tree mortality year (50mm) | 0.60 | 0.43 – 0.86 | 0.005 |

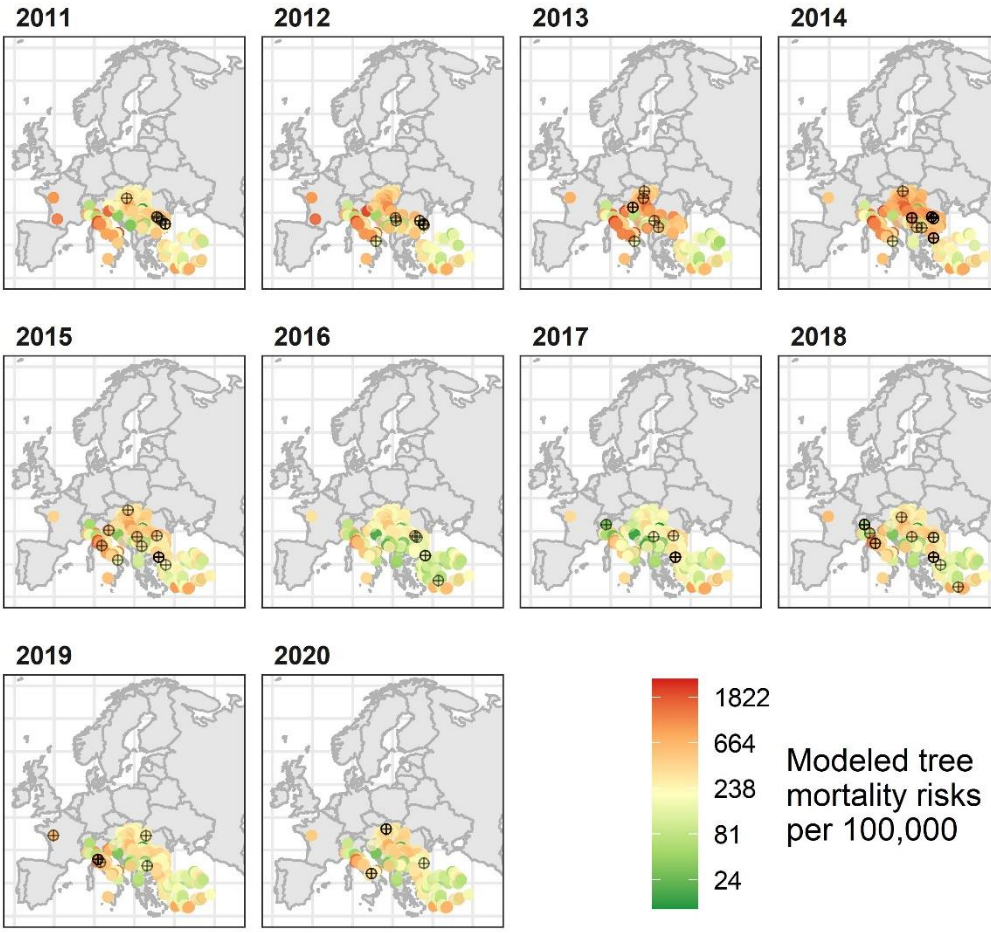


**Fig S7** Spatial evaluation of modelled tree mortality for Austrian oak for all analyzed ICP Forests plots on a logarithmic color scale. Black points indicate observations of dead trees (n=101)

European hornbeam

**Table S7.** Logistic regression model estimates for European hornbeam; irregular aged trees excluded due to lack of tree mortalities in this group.

| Variable | Odds ratio | 95% Confidence interval | P-value |
|---|---|---|---|
| Mean stand age (Reference 41 -60) | | | 0.0003 |
| 61 – 80 | 1.32 | 0.77 – 2.28 | |
| 81 – 100 | 1.25 | 0.64 – 2.39 | |
| 101 - 120 | 1.46 | 0.68 – 2.97 | |
| >120 | 4.76 | 2.49 – 8.98 | |
| Temperature difference between average temperature of the coldest and the warmest month (continentality) 30-years average (°C) | 1.46 | 1.26 – 1.73 | < 0.0001 |
| Total precipitation in vegetation period 30-years average (50mm) | 0.83 | 0.70 – 0.97 | 0.02 |
| Total precipitation in winter anomaly average of the reported tree mortality year and the previous year (50mm) | 1.78 | 1.25 – 2.49 | 0.002 |

| | | | |
|---|---|---|---|
| Average temperature of the warmest month previous year to reported tree mortality (°C) | 1.54 | 1.15 – 2.09 | 0.003 |

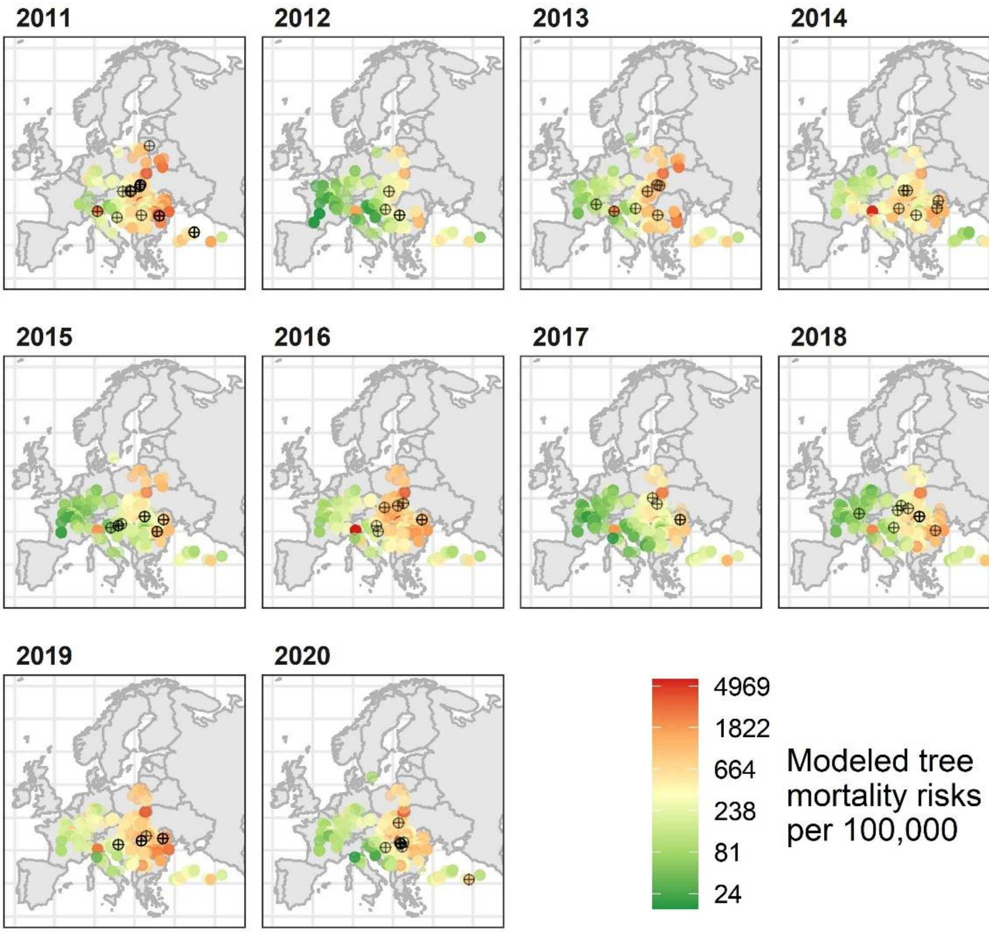


**Fig S8** Spatial evaluation of modelled tree mortality for European hornbeam for all analyzed ICP Forests plots on a logarithmic color scale. Black points indicate observations of dead trees (n=103)

Pedunculate oak

**Table S8.** Logistic regression model estimates for Pedunculate oak.

| **Variable** | **Odds ratio** | **95% Confidence interval** | **P-value** |
|---|---|---|---|
| Average temperature in summer 30-years average (°C) | 1.28 | 1.19 – 1.39 | < 0.0001 |
| Total precipitation in summer 30-years average (50mm) | 1.34 | 1.19 – 1.48 | < 0.0001 |
| Average temperature of the warmest month anomaly average of the 3 previous year to reported tree mortality (°C) | 1.97 | 1.42 – 2.73 | < 0.0001 |
| Average temperature of the coldest month anomaly average of the reported tree mortality year (°C) | 1.19 | 1.09 – 1.30 | < 0.0001 |
| Total precipitation in winter anomaly average of 2 previous years to the reported tree mortality year (50mm) | 1.86 | 1.39 – 2.46 | < 0.0001 |
| Interaction between average temperature of the coldest month anomaly average of the reported tree mortality year and total | 0.78 | 0.66 – 0.91 | 0.002 |

precipitation in winter anomaly average of 2 previous years to the reported tree mortality year (°C * 50mm)

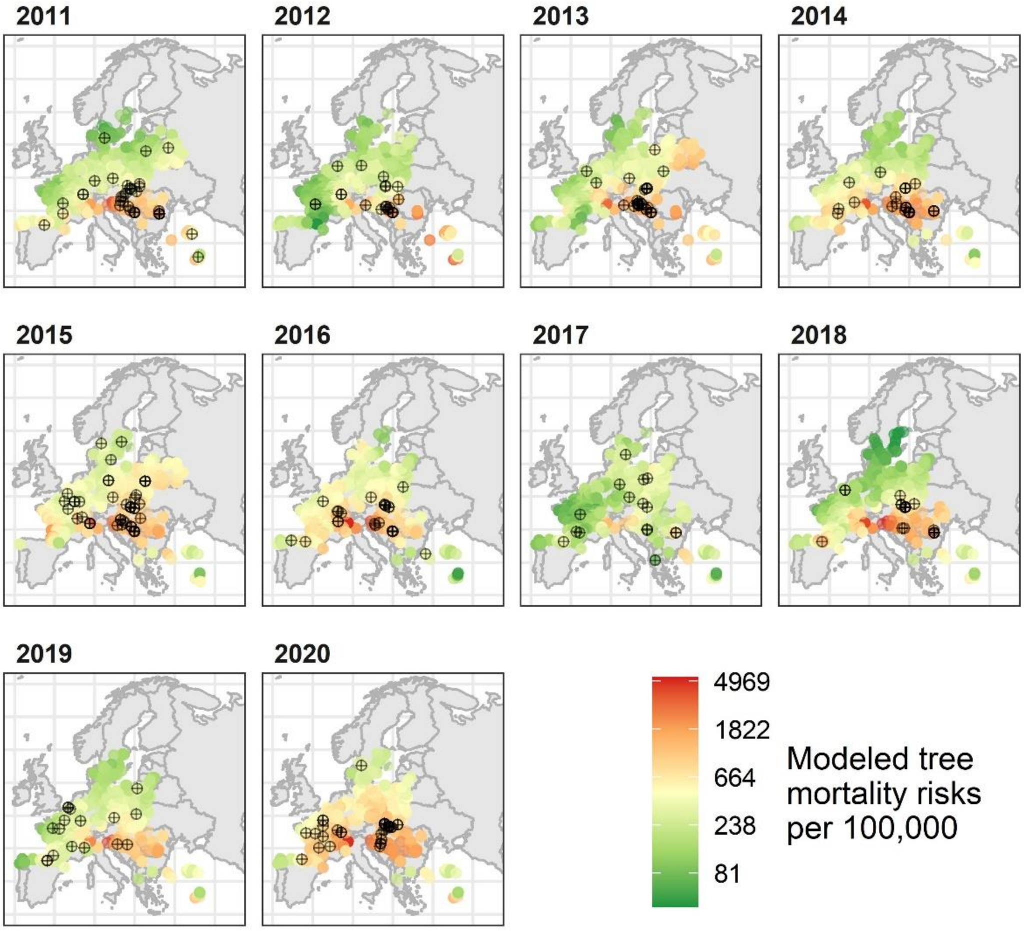


**Fig S9** Spatial evaluation of modelled tree mortality for Pedunculate oaks for all analyzed ICP Forests plots on a logarithmic color scale. Black points indicate observations of dead trees (n=277)

Sessile oak

**Table S9.** Logistic regression model estimates for Sessile oak.

| **Variable** | **Odds ratio** | **95% Confidence interval** | **P-value** |
|---|---|---|---|
| Mean stand age (Reference 41 -60) | | | < 0.0001 |
| 61 – 80 | 0.73 | 0.50 – 1.05 | |
| 81 – 100 | 0.39 | 0.25 – 0.57 | |
| 101 - 120 | 0.37 | 0.20 – 0.63 | |
| >120 | 0.48 | 0.25 – 0.84 | |
| Irregular stands | 0.43 | 0.21 – 0.82 | |
| Aspect (Reference Flat) | | | 0.0001 |
| North | 2.42 | 1.46 – 4.06 | |
| North-east | 1.29 | 0.71 – 2.31 | |
| East | 1.48 | 0.87 – 2.54 | |
| South-east | 1.50 | 0.83 – 2.70 | |
| South | 0.74 | 0.34 – 1.50 | |

| | | | |
|---|---|---|---|
| South-west | 0.78 | 0.42 – 1.42 | |
| West | 1.11 | 0.59 – 2.02 | |
| North-west | 0.93 | 0.44 – 1.85 | |
| Temperature difference between average temperature of the coldest and the warmest month (continentality) 30-years average (°C) | 0.59 | 0.41 – 0.82 | 0.002 |
| Annual total precipitation 30-years average (50mm) | 0.31 | 0.19 – 0.49 | < 0.0001 |
| Interaction between temperature difference between average temperature of the coldest and the warmest month 30-years average and annual total precipitation 30-years average (°C * 50mm) | 1.06 | 1.04 – 1.09 | < 0.0001 |
| Average temperature in spring anomaly average of the reported tree mortality year and 2 previous years (°C) | 0.44 | 0.29 – 0.67 | 0.0001 |
| Average temperature in winter anomaly average of the reported tree mortality year and 3 previous years (°C) | 0.58 | 0.47 – 0.70 | < 0.0001 |
| Total precipitation in summer anomaly average of 2 previous years to the reported tree mortality year (50mm) | 0.98 | 0.80 – 1.19 | 0.8 |
| Interaction between average temperature in winter anomaly average of the reported tree mortality year and 3 previous years and total precipitation in summer anomaly average of 2 previous years to the reported tree mortality year (°C * 50mm) | 0.64 | 0.50 – 0.83 | 0.001 |

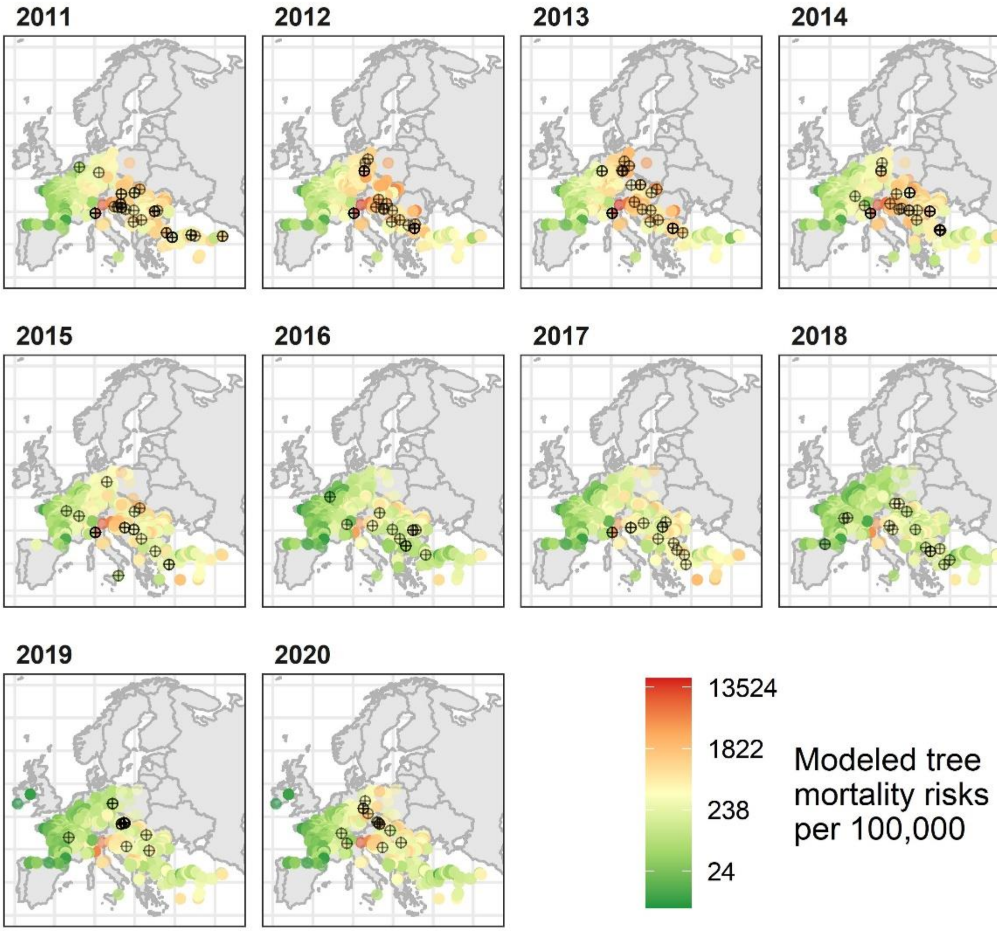

**Fig S10** Spatial evaluation of modelled tree mortality for Sessile oaks for all analyzed ICP Forests plots on a logarithmic color scale. Black points indicate observations of dead trees (n=232)

**Table S10.** Mortality classification.

| Description | Category | Mortality |
|---|---|---|
| Utilization for biotic reasons, e.g. insect damage | Tree has been cut and removed, only its stump has been left | Yes |
| Utilization for abiotic reasons, e.g. windthrow | Tree has been cut and removed, only its stump has been left | Yes |
| Biotic reasons, e.g. bark beetle attack | Standing dead tree | Yes |
| Abiotic reasons, e.g. drought, lightning | Standing dead tree | Yes |
| Unknown cause of death | Standing dead tree | Yes |
| Tree alive and measurable (new, note this is different than a missing value) | Tree alive | No |
| Tree alive, in current and previous inventory | Tree alive | No |
| New alive tree (ingrowth) | Tree alive | No |
| Alive tree (present but not assessed in previous inventory, incl. replacement trees) | Tree alive | No |
